# A Multiscale Spatial Modeling Framework for the Germinal Center Response

**DOI:** 10.1101/2024.01.26.577491

**Authors:** Derek P. Mu, Christopher D. Scharer, Norbert E. Kaminski, Qiang Zhang

**Affiliations:** Montgomery Blair High School, Silver Spring, MD 20901, USA; Department of Microbiology and Immunology, School of Medicine, Emory University, Atlanta, GA 30322, USA; Department of Pharmacology & Toxicology, Institute for Integrative Toxicology, Michigan State University, East Lansing, Michigan 48824, USA; Gangarosa Department of Environmental Health, Rollins School of Public Health, Emory University, Atlanta, GA 30322, USA

**Author notes:** Corresponding author: Qiang Zhang (ORCID: 0000-0002-8678-7013) Gangarosa Department of Environmental Health Rollins School of Public Health Emory University 1518 Clifton Rd NE Atlanta, GA 30322, USA.

**Keywords:** B cells, germinal center, dark zone, light zone, affinity maturation, proliferative burst, chemotaxis

## Abstract

The germinal center response or reaction (GCR) is a hallmark event of adaptive humoral immunity. Unfolding in the B cell follicles of the secondary lymph organs, a GC culminates in the production of high-affinity antibody-secreting plasma cells along with memory B cells. By interacting with follicular dendritic cells (FDC) and T follicular helper (Tfh) cells, GC B cells exhibit complex spatiotemporal dynamics. Driving the B cell dynamics are the intracellular signal transduction and gene regulatory network that responds to cell surface signaling molecules, cytokines, and chemokines. As our knowledge of the GC continues to expand in depth and in scope, mathematical modeling has become an important tool to help disentangle the intricacy of the GCR and inform novel mechanistic and clinical insights. While the GC has been modeled at different granularities, a multiscale spatial simulation framework – integrating molecular, cellular, and tissue-level responses – is still rare. Here, we report our recent progress toward this end with a hybrid stochastic GC framework developed on the Cellular Potts Model-based CompuCell3D platform. Tellurium is used to simulate the B cell intracellular molecular network comprising NF-κB, FOXO1, MYC, AP4, CXCR4, and BLIMP1 that responds to B cell receptor (BCR) and CD40-mediated signaling. The molecular outputs of the network drive the spatiotemporal behaviors of B cells, including cyclic migration between the dark zone (DZ) and light zone (LZ) via chemotaxis; clonal proliferative bursts, somatic hypermutation, and DNA damage-induced apoptosis in the DZ; and positive selection, apoptosis via a death timer, and emergence of plasma cells in the LZ. Our simulations are able to recapitulate key molecular, cellular, and morphological GC events including B cell population growth, affinity maturation, and clonal dominance. This novel modeling framework provides an open-source, customizable, and multiscale virtual GC simulation platform that enables qualitative and quantitative *in silico* investigations of a range of mechanic and applied research questions in future.

## Introduction

The adaptive humoral immune response is a vital component of host defense, where B cells terminally differentiate into plasma cells (PCs) that secrete antibodies specifically recognizing and neutralizing the invading foreign antigens. The B cell responses can be broadly classified into two types: T cell-dependent and independent, depending on whether helper T (Th) cells are involved (Nutt et al. 2015). In the T cell-independent response, naive B cells are activated directly via toll-like receptors (TLR) recognizing pathogen components such as lipopolysaccharide (LPS) or CpG DNA or via B cell receptors (BCR), without the assistance of Th cells (Fagarasan and Honjo 2000, Allman et al. 2019). The response is launched quickly and can occur within a few days of initial infection. Upon activation, B cells undergo clonal proliferation and differentiate into PCs, which secrete pentameric IgM molecules. While these antibodies provide initial protection, they are often polyclonal, not highly specific, and the IgM-secreting PCs are short-lived, thus unable to provide long-term immunity. In contrast, the T cell-dependent B cell response takes longer to develop, but through affinity maturation and class switch recombination (CSR) it can produce long-lived PCs that can provide life-long immunity with high-affinity IgG or other non-IgM antibody classes (Parker 1993). Additionally, memory B cells are generated during the primary response, which can quickly launch a secondary antibody response upon subsequent exposure to the same antigens (Inoue and Kurosaki 2024).

T cell-dependent B cell activation takes place primarily in the germinal centers (GC), which are specialized, often transient, microstructures formed in the B cell follicles of secondary lymphoid tissues such as lymph nodes and spleen in response to infection or immunization (Mesin et al. 2016, Stebegg et al. 2018, Young and Brink 2021, Victora and Nussenzweig 2022). GC B cells exhibit unique spatiotemporal dynamics (Allen et al. 2007a, Victora et al. 2010). A GC is polarized, containing two distinct, physically separated zones: the dark zone (DZ) and the light zone (LZ). In the DZ, B cells undergo clonal proliferative bursts, during which somatic hypermutation (SHM) occurs. During SHM, the hypervariable regions of the genes encoding the immunoglobulin heavy chains and light chains are point-mutated by activation-induced cytidine deaminase (AID) at a high rate (Methot and Di Noia 2017). As a result, the BCR affinities of the participating B cell clones for the invading antigen are modified and diversified. During SHM those B cells incurring damaging mutations that prevent normal assembly of surface BCRs are killed via apoptosis in the DZ (Mayer et al. 2017, Stewart et al. 2018). After exiting the cell cycle following a proliferative burst, B cells migrate from the DZ to LZ under the chemoattractant force by CXCL13 secreted by follicular dendritic cells (FDCs) in the LZ (Allen et al. 2004, Cosgrove et al. 2020).

In the LZ, two main cell types participate in the positive selection of B cell clones harboring immunoglobulin (Ig) gene variants encoding relatively high-affinity antibodies: the residential FDCs and CD4^+^ T follicular helper (Tfh) cells. These two cell types coordinate to provide key molecular signals for B cell activation, survival, DZ re-entry, proliferation, and differentiation (Vinuesa et al. 2010, Luo et al. 2018, Crotty 2019). FDCs are antigen-presenting cells (APCs), which previously encountered and engulfed pathogens and present the antigen epitopes on their cell surface. When B cells first encounter FDCs, their BCRs are activated by the surface antigens of FDCs. B cells then internalize the antigen-BCR complex and present the antigen epitopes on their own surface through major histocompatibility complex (MHC) II molecules to form peptide-MHCII complex (pMHCII). The density of pMHCII on the cell surface is proportional to the BCR affinity. Stronger BCR signaling also leads to higher PI3K-AKT-FOXO1 signal transduction (Hinman et al. 2007, Sander et al. 2015, Luo et al. 2018). When the B cells subsequently encounter Tfh cells, a complex mutual interaction occurs between the two cell types (Ise et al. 2018, Crotty 2019, Mintz and Cyster 2020). Tfh cells are activated via T cell receptors (TCR) liganded by pMHCII of the B cells, as well as by other cell surface signaling molecules such as inducible co-stimulator ligand (ICOSL) (Liu et al. 2015). Activated Tfh cells in turn express surface CD40L which reciprocally activates B cells together with several secretory cytokines including interleukins (IL) 4, 10, and 21 (Reinhardt et al. 2009, Xin et al. 2018, Quast et al. 2022). CD40 signaling leads to NF-κB activation, increasing the chance of survival of B cells. In the presence of downregulated FOXO1, NF-κB elicits transient MYC activation that initiates the cell cycle (Luo et al. 2018). Only a small fraction of B cells is positively selected, which express CXCR4, the receptor for chemokine CXCL12, and migrate back to DZ where they undergo further proliferative bursts and SHM (Allen et al. 2004, Sander et al. 2015). Those B cells with weaker BCR affinity are more likely to undergo apoptosis in the LZ, as well as B cells that do not have a chance to encounter Tfh cells in the LZ. As a result of combined action of proliferation and SHM in the DZ and positive selection in the LZ, the overall BCR affinity of the GC B cell population for the antigen continues to improve. After many rounds of DZ-LZ cycles, a small fraction of B cells are affinity-matured and exit the GC as either long-lived antibody-secreting PCs or memory B cells.

The GC plays a critical role in the generation of long-term protective immunity, and this is relevant in both the context of natural infection and vaccination for infectious diseases such as COVID-19 (Laidlaw and Ellebedy 2022). If the GC is compromised or cannot be sufficiently induced due to genetic alterations, increased susceptibility to bacterial and viral infections will result. On the other hand, unintentional recognition of self-antigens and induction of GC can lead to autoimmune diseases such as systemic lupus erythematosus (Woods et al. 2015). Dysregulated B cell proliferation in GC can lead to lymphoma or other B cells-related leukemia (Mlynarczyk et al. 2019). GC also plays a role in antibody-mediated rejection of transplanted organs (Chong 2020). In addition, many environmental contaminants are immunotoxicants, some of which can suppress B cell activation and the humoral immune response, leading to increased susceptibility to infectious disease and cancer (Germolec et al. 2022). Therefore, a full mechanistic understanding of the complexity of GC is crucially important for sustaining immune integrity and preventing or alleviating many pathological conditions.

Computational modeling has played a long-standing role in dissecting and understanding the complex dynamics of GC immune responses (Meyer-Hermann et al. 2009, Meyer-Hermann et al. 2012). The GC involves an elaborate interplay between many cell types, signaling molecules, transcription factors, and actuator genes (Verstegen et al. 2021). Key signal transduction and gene regulatory networks underpin the spatiotemporal dynamics of GC B cells and are crucial for the positive selection and ultimate formation of high-affinity PCs. Although there have been many efforts simulating the cellular dynamics and affinity maturation of GCs, cross-scale modeling that integrates molecular, cellular, and tissue-level actions in a spatial context has only begun to emerge recently and thus is still rare (Merino Tejero et al. 2021a, Merino Tejero et al. 2021b, Merino Tejero et al. 2022). In this study, we presented a novel multiscale mathematical modeling framework of the GC developed in the CompuCell3D simulation environment that integrates the molecular network and spatiotemporal behaviors of GC B cells. The modeling framework provides an open-source, customizable, multiscale virtual GC platform that enables future *in silico* investigations of a range of questions both quantitatively and quantitatively, including B cell population turnover, BCR mutation rate, death timer, proliferative burst size, availability of Tfh cells, and effects of genetic and chemical perturbations.

## Methods

### 1. Model structure

#### 1.1 Cell types, cellular events, and interactions

Four cell types are modeled in the framework: CXCL12-expressing reticular cells (CRCs) located in DZ, FDCs and Tfh cells in LZ, and B cells cycling between the DZ and LZ. For simplicity, CRC, FDC, and Tfh cells are treated as stationary. Key cellular events of B cells captured in the model include: (i) B cell volume growth, division, SHM, and apoptosis in the DZ; (ii) DZ-to-LZ B cell migration and simultaneous initiation of a cell death timer; (iii) interaction of B cells with FDCs in the LZ to determine BCR antigen affinity, interaction of B cells with Tfh cells in the LZ to make probabilistic decisions based on pMHC density on positive selection, survival, initiation of cell growth, and DZ re-entry of positively selected B cells, and death timer-triggered apoptosis of LZ B cells not positively selected.

#### 1.2 Molecular events in B cells

The above cellular events are driven by an intracellular molecular network in B cells that responds to diffusive chemoattractants and signaling molecules from FDCs and Tfh cells. For simplicity, the following molecular species and regulatory events are included in the model (Fig. 1). A DZ-to-LZ descending gradient of chemoattractant CXCL12 is established by CRCs in DZ (Bannard et al. 2013, Rodda et al. 2015), and an opposite gradient of chemoattractant CXCL13 is established by FDCs in LZ (Wang et al. 2011, Cosgrove et al. 2020). In LZ, the contact of a B cell with an FDC will trigger BCR-mediated signal transduction, which leads to several signaling events in the modeled B cell: (1) re-expression on B cell surface of pMHCII, the density of which depends on BCR affinity for the antigen, (2) transient activation of AKT and downregulation of FOXO1, the extent of which depends on BCR affinity (Hinman et al. 2007, Sander et al. 2015, Luo et al. 2018), (3) once the BCR affinity reaches a threshold, a switch-like activation of NF-κB subtype RelA is triggered (Shinohara et al. 2014, Michida et al. 2020, Wibisana et al. 2022), which in turn induces BLIMP1 (Heise et al. 2014, Roy et al. 2019), leading to terminal B cell differentiation into antibody-secreting PCs.

**Figure 1.**
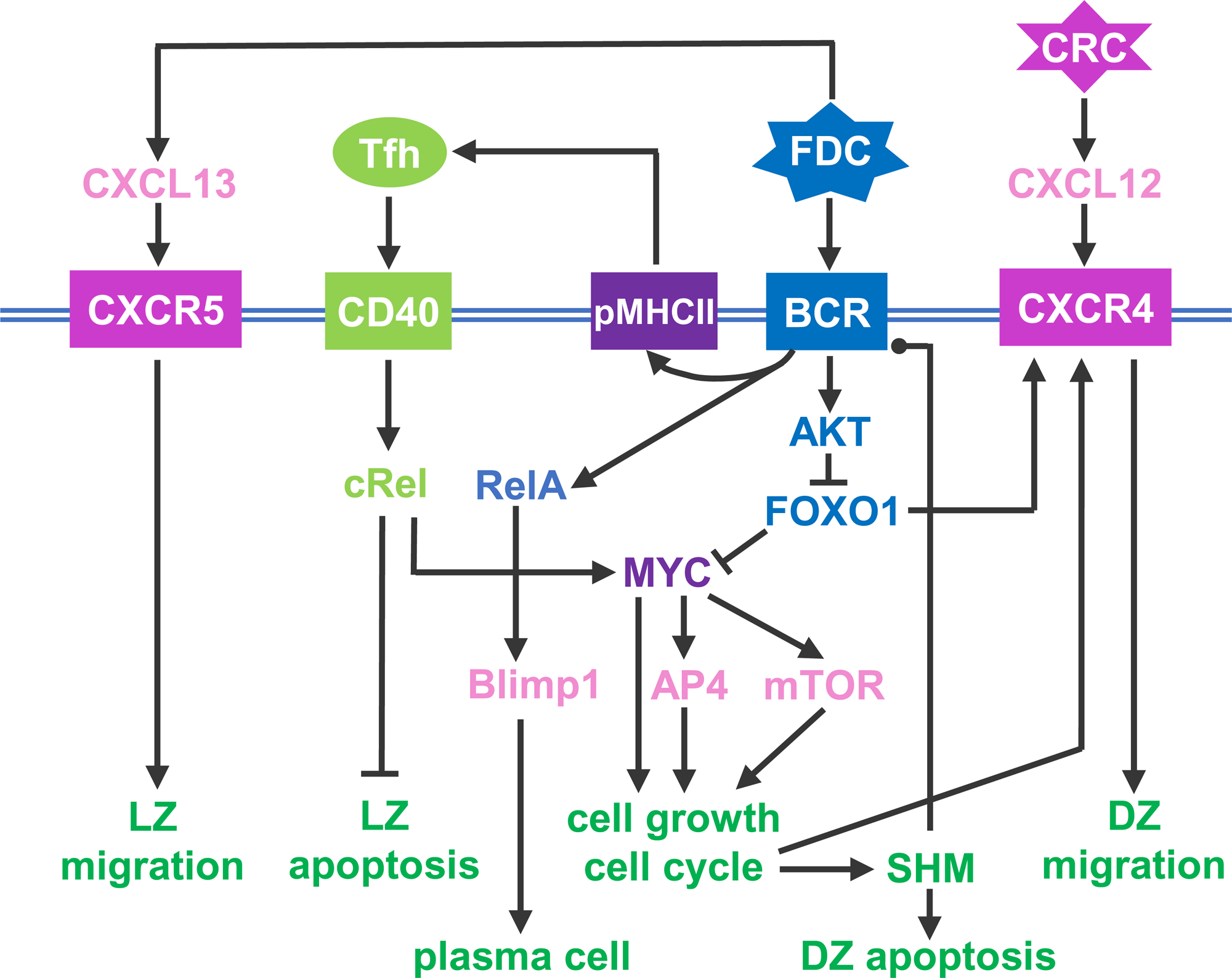
Schematic illustration of a simplified intracellular molecular network of GC B cells and different B cell outcomes driven by key molecules as indicated. Pointed arrowhead: stimulation/activation, blunted arrowhead: inhibition, and dotted arrow head: regulation in either direction.

If a B cell expressing pMHCII encounters a Tfh cell, the B cell is stimulated via CD40 signaling which activates another NF-κB subtype, cRel. cRel activation leads to at least two molecular signaling events. It activates BclxL which inhibits apoptosis, thus terminating the death timer (Zarnegar et al. 2004). With FOXO1 still downregulated, cRel also induces the expression of MYC (Luo et al. 2018). Upregulation of MYC triggers commitment to cell growth and initiates the cell cycle (Dominguez-Sola et al. 2012). MYC also activates AP4, which sustains B cell growth and division burst (Chou et al. 2016). With the B cell committed to growth and proliferation, as FOXO1 is re-expressed, CXCR4 is induced (Dominguez-Sola et al. 2015, Sander et al. 2015). As a result, the B cell migrates towards the DZ in response to the CXCL12 gradient. In the DZ, shortly after each cell division, each of the two daughter cells incurs an independent point mutation of BCR which alters its affinity for the antigen with some probabilities (Faili et al. 2002, Sharbeen et al. 2012, Wang et al. 2017). The probability of a damaging mutation is encoded such that a fraction of B cell progeny dies by apoptosis in the DZ (Mayer et al. 2017, Stewart et al. 2018). The surviving B cells will continue to grow and divide as long as AP4 remains above a threshold (Chou et al. 2016). When AP4 drops below the threshold, the B cells exit the cell cycle followed shortly by downregulation of CXCR4 (Weber 2018). With CXCR5 constitutively expressed (Allen et al. 2004, Victora et al. 2010), the B cells will be pulled by the CXCL13 gradient field into the LZ, repeating the DZ-LZ cycle.

#### 1.3 Model assumptions and simplification

The molecular and cellular events involved in the GC are complex. Here, for the purpose of modeling, several assumptions and simplifications are made.

1. The mutual activation between B cells and Tfh cells is simplified to pMHCII density-dependent CD40-cRel activation, as described above.
2. Many factors involved in GC, including BCL6, BACH2, and IRF4, are not explicitly considered.
3. Initial migration of B cells from the T/B cell border towards the LZ is not considered and neither is the Tfh cell migration into the LZ. CSR is thus not included as it is believed to occur primarily during pre-GC formation (Roco et al. 2019).
4. For B cells returning to the DZ, a delay variable is introduced before the cell growth for the first cell cycle is initiated to reduce the chance of cell division in the LZ.
5. The GC exit of PCs is modelled as deleting these cells from the simulation once they emerge.
6. Formation of memory B cells is not considered.

### 2. Construction of the computational model in CompuCell3D

The GC model was constructed and simulated as a hybrid, agent-based stochastic model in CompuCell3D. CompuCell3D provides a flexible and customizable platform for simulating multi-cellular behaviors and interactions based on the Glazier-Graner-Hogeweg approach (Swat et al. 2012). The Cellular Potts Model module in CompuCell3D was employed to simulate the physical properties and movements of individual B cells (Graner and Glazier 1992), while the molecular network operating in each individual B cell, as depicted in Fig. 1, was simulated by using the Gillespie’s stochastic algorithm implemented in Tellurium conforming to the Antimony notation (Gillespie 1977, Choi et al. 2018). The CompuCell3D model consists of four files: an XML file, a Potts initialization file (PIF), and two Python script files. The XML file contains various “Plugins” and “Steppables” that define some default Potts model parameter values. The *Chemotaxis* plugin defines CXCL12 and CXCL13 as the chemoattractants, and the *DiffusionSolverFE* steppable designates that CXCL12 and CXCL13 are secreted by CRC and FDC respectively and specifies the parameter values for secretion, diffusion, and decay in the Medium. The PIF file contains the initial coordinates of medium and cells where applicable. The steppable Python file contains the script that defines several steppable classes, including *GCR_Steppable*, *MitosisSteppable*, *BCell_GRNSteppable*, and *VisualizationSteppable*, and the Tellurium model. The main Python file contains the script that imports and registers all the steppables and runs the simulation.

#### 2.1 Initialization

The model is initialized in the “start” section of *GCR_Steppable*, including the generation of CRCs, FDCs, Tfh cells, and seeding B cells. Each B cell is assigned a Tellurium molecular network model named as *BCellNetwork*.

#### 2.2 Cell-cell contact and probabilistic decision-making

To capture the physical contact between B cells, FDCs, and Tfh cells, the *NeighborTracker* and *PixelTracker* plugins are employed to identify the neighboring cells of each B cell. All neighboring cells of a B cell at a given moment are first identified by utilizing the *get_cell_neighbor_data_list()* function, then the specific cell type of each neighboring cell is identified by utilizing the *neighbor_count_by_type()* function. Once a contact with an FDC is identified, the pMHCII level of the B cell is set proportional to its antigen-specific BCR affinity. Upon subsequent contact with a Tfh cell, the B cell can be positively selected based on a probability that is proportional to the pMHCII level. For the positively selected cell, the running death timer is terminated, cell cycle is committed, and DZ re-entry is initiated.

#### 2.3 SHM and probability of BCR affinity alteration

Each of the two daughter cells of a dividing B cell has a probability of 0.3 to produce a damaging mutation that will result in cell death in the DZ. For the daughter cell that does not incur a damaging mutation, an SHM can either increase, decrease, or does not change the BCR affinity, each with a probability of 1/3. The increment or decrement of the affinity alteration can be either 0.25 or 0.5 with equal probability. In general, the BCR affinity ranges between 0-10, but can be higher.

### 3. Simulation data collection, storage, analysis, and model sharing

Variables of each B cell are saved in a plain text file which is updated every 15 Monte Carlo steps (*mcs*). Saved variables include cell generation, mother ID, cell ID, BCR affinity, cell volume, X, Y, and Z-coordinates, and the molecular species levels in the Tellurium model, etc. The file is named in the format of “Generation_mother ID_cell ID.txt”. Analyzing the simulation results saved in the txt files was conducted in a separate Python script. The CompuCell3D model files, which contain the model parameter values, are available as Supplemental Material.

## Results

### 1. Morphology of a simulated GC

The morphological results of a representative simulation of the GC model are shown in Fig. 2. The simulated space dimension is 250×200×11 pixels, which can be considered as 250×200×11 in µm in real space, to represent a slice of the GC to save computational time. The 2-D projection of the instantiated non-B cells on the X-Y plane is indicated in Fig. 2A, while along the Z dimension, these cells are distributed randomly in 3 of the 11 layers (results not shown). There are 45 CRCs distributed on the left half of the field, which becomes the future DZ, and 45 FDCs and 36 Tfh cells distributed on the right half of the field, which becomes the future LZ. Tfh cells are located next to FDCs, reflecting the notion that they also express CXCR5 and are thus drawn to the LZ by chemoattractant CXCL13 (Breitfeld et al. 2000, Kim et al. 2001). Not all FDCs are surrounded by Tfh cells, mimicking the situation that the availability of Tfh cells is a limiting factor in the positive selection of LZ B cells (Meyer-Hermann et al. 2006, Allen et al. 2007b, Meyer-Hermann 2007, Victora et al. 2010). CRCs and FDCs secrete CXCL12 and CXCL13 respectively, establishing two opposing chemoattractant fields and thus the polarity of the GC (Fig. 2B and 2C).

**Figure 2.**
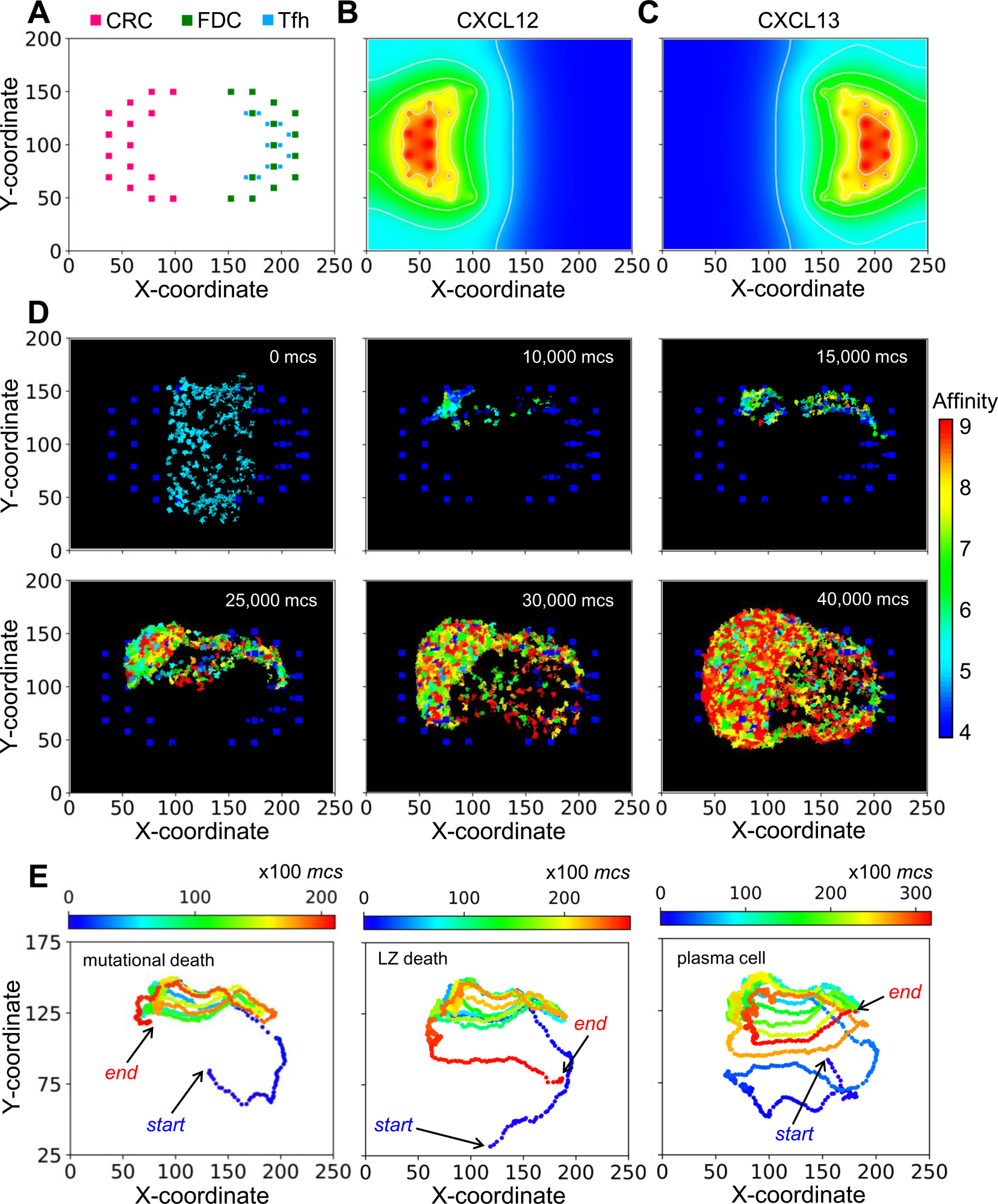
Morphology of a representative simulated GC. **(A)** 2-D distributions of CRCs, FDCs, and Tfh cells in the DZ (left) and LZ (right) with X-Y coordinates indicated. **(B-C)** Concentration gradients of chemoattractants CXCL12 and CXCL13 respectively. White contour lines: isolines of equal concentrations. **(D)** Snapshots of simulated GC at various *mcs* indicated. Colors of B cells denote BCR/antibody affinity as indicated by the colormap. **(E)** 2-D trajectories of B cells in three select lineages leading respectively to damaging mutation-induced apoptosis in DZ (left panel), death timer-triggered apoptosis in the LZ for not positively selected (mid panel), and emergence of a PC (right panel). Dot color denotes *mcs* time as indicated by the colormap.

With the layout of residential cells and chemoattractant fields as above, a simulated GC that succeeds in producing B cells of high BCR/antibody affinities is shown as snapshots in Fig. 2D and in Supplemental Video S1. For this simulation, the GC starts with 200 B cells (clones) of intermediate affinity of 5 (indicated by the color of the cells) between the DZ and LZ. Over a period of 40,000 Monte Carlo steps (*mcs*, where 100 mcs can be regarded approximately as 1 hour in real time), both the number of total GC B cells and the fraction of high-affinity B cells increase, indicating successful GC population growth and affinity maturation. The 2-D trajectories of 3 select B cell lineage branches leading to different cell fates are shown in Fig. 2E. These trajectories cycle between the DZ and LZ for multiple rounds. The first trajectory ends with cell death in the DZ due to damaging mutation during mitosis (left panel), the second trajectory also ends with cell death but in the LZ due to death timer (mid panel), and the last trajectory ends with differentiation into a PC in the LZ (right panel).

### 2. Cellular events of GC B cells

#### 2.1 B cell population dynamics

In this section we performed an in-depth quantitative analysis of the GC B cell population dynamics with respect to time, location, and cell fates. For the GC simulation presented in Fig. 2, the total number of B cells (*N_Tot_*) increases rapidly from the initial 200 over a period of 40,000 *mcs*, then approaches a steady-state size of about 2,500 cells through 72,000 *mcs* (equivalent to 30 days) without considering GC termination (Fig. 3A). The growth of the B cell population is not smooth – it proceeds in an uneven fashion due to random births and deaths occurring simultaneously. In the early stage of the GC, the numbers of B cells in the DZ (*N_DZ_*) and LZ (*N_LZ_*) alternate in anti-phase, resulting from cyclic cell migration in unison between the two zones (Video S1). In the late stage, *N_DZ_* is persistently greater than *N_LZ_*with the *N_DZ_*:*N_LZ_* ratio stabilizing near 3:1 (Fig. 3A). The evolving B cell population in the GC is highly dynamic with constant turnover through several processes. Specifically, B cells (i) are born in the DZ out of proliferative bursts of clonal expansion, (ii) are cleared from the GC via apoptosis due to damaging BCR mutations in the DZ and, if not positively selected, in the LZ, and (iii) exit the GC as PCs. We next quantified the birth and death events.

**Figure 3.**
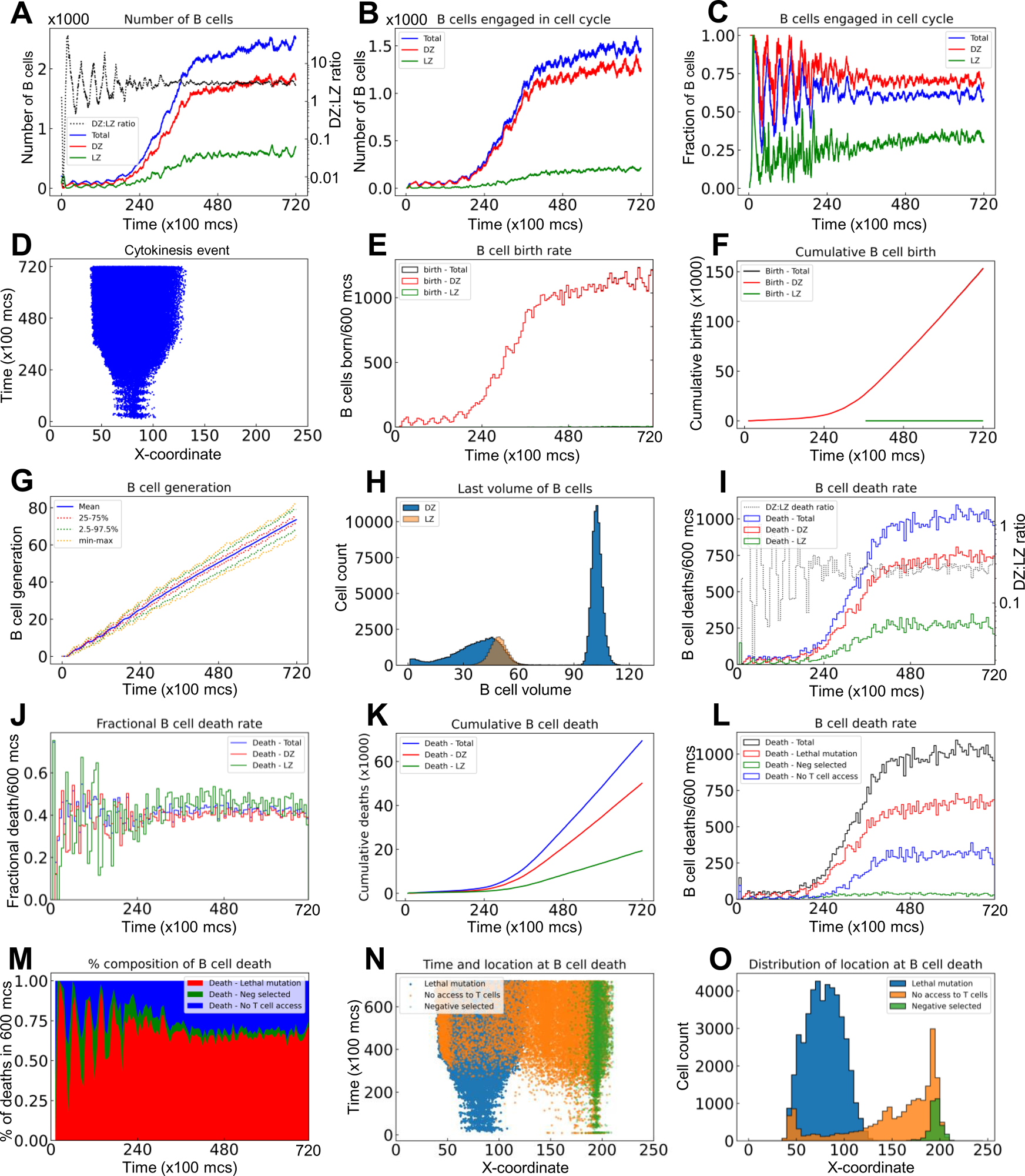
Quantitative analyses of B cell population dynamics in a simulated GC. **(A)** Numbers of B cells in the DZ, LZ, and total, and DZ:LZ birth ratio as indicated at a given time. **(B)** Numbers of B cells engaged in cell cycle in DZ, LZ, and total as indicated. **(C)** Fractions of B cells in the DZ, LZ, and total that are engaged in cell cycle. **(D)** The X-coordinate and time at which cytokinesis events occur. **(E)** Numbers of B cells born every 600 *mcs* in the DZ, LZ, and total as indicated. **(F)** Numbers of cumulative cell births in the DZ, LZ, and total as indicated. **(G)** Mean, interquartile, 2.5-97.5^th^ percentile, minimum and maximum generations of GC B cells as indicated. **(H)** Distributions of last volumes of B cells in the DZ and LZ as indicated before they divide, die, or differentiate into PCs. **(I)** Numbers of B cell deaths in every 600 *mcs* in the DZ, LZ, and total, and DZ:LZ death ratio as indicated. **(J)** Fractions of B cells in the DZ, LZ, and total that die in every 600 *mcs* as indicated. **(K)** Numbers of cumulative cell deaths in the DZ, LZ, and total as indicated. **(L)** Numbers of B cell deaths in every 600 *mcs* in total, due to damaging BCR (lethal) mutation, not being positively (Neg) selected after contacting Tfh cells, or no access to Tfh cells as indicated. **(M)** Percentage composition of B cell deaths in every 600 *mcs* due to lethal mutation, negative selection, or no access to Tfh cells as indicated. **(N)** The X-coordinate and time at which cell deaths occur. **(O)** Distributions of X-coordinate at which cell deaths occur due to lethal mutation, negative selection, or no access to Tfh cells as indicated.

##### 2.1.1 B cell birth

The number of B cells engaged in cell cycle increases over time and these cells are predominantly in the DZ (Fig. 3B). There are a small number of cell cycle-engaged B cells in the LZ, representing positively selected B cells that just initiate the cell cycle without much growing yet. Approaching the steady state, nearly 70% and 30% of DZ and LZ B cells, respectively, are engaged in the cell cycle, while on average 65% of the overall B cell population is in the cell cycle (Fig. 3C). B cells are born predominantly in the DZ with only a negligible number of births in the LZ (Fig. 3D). The absolute birth rate increases over time approaching about 1100 births per 600 *mcs* (Fig. 3E). Cumulative births reach 150K in the entire 72,000 *mcs* period when two birth events are registered for each cell division (Fig. 3F). The mean cell generation increases almost linearly with time while the variability also progressively increases as more B cells are born (Fig. 3G). The DZ B cell volumes exhibit a biphasic distribution, reflecting that these cells are actively engaged in growth and division (Fig. 3G). In contrast, the LZ B cells exhibit a very narrow volume distribution consistent with the notion that they are mostly non-proliferating centrocytes.

##### 2.1.2 B cell death

As the GC B cell population grows, the number of cell deaths increases and then approaches a steady state, where the total death rate is about 1000 deaths per 600 *mcs* (Fig. 3I). Although at the early time the DZ:LZ death rate ratio fluctuates dramatically as a result of randomness due to small numbers of cell deaths and DZ-LZ migration, the ratio stabilizes at about 2.5:1 at later time. The steady-state death turnover rate of the overall B cell population is slightly above 40% in 600 *mcs*, and the turnover rates in both zones are similar (Fig. 3J). Cumulatively, there are nearly 70,000 cell deaths, among which 70% occurred in the DZ and 30% in the LZ (Fig. 3K).

Further analysis showed that different types of cell deaths occur at different rates (Fig. 3L). B cell death due to damaging BCR mutations occurs most often, comprising 65% of total deaths at steady state, followed by cell death due to no access to Tfh cells at 30%, while cell death for not being positively selected even after contacting Tfh cells (labelled as Neg selected) is only a small fraction of all death events (Fig. 3M). When a B cell cannot access Tfh cells, positive selection decisions cannot be made in time and the B cell will die when the default death timer goes off. This type of death is only limited to B cells in the LZ initially, but as the GC population grows such that the space becomes more compact and thus crowded, such death also expands to the DZ when some of the B cells exiting the cell cycle do not have enough time to migrate through the densely populated DZ (Fig. 3N); however, these deaths in the DZ are only a small fraction (Fig. 3O).

#### 2.2 Affinity maturation and clonal dominance

We next characterized the evolution of the BCR antigen affinities in the simulated GC. With all the 200 seeder B cells starting with an intermediate BCR affinity of 5 in this simulation, their clonal affinities initially drift to both higher and lower levels (Fig. 4A). However, the mean affinity increases progressively in a winding manner and then reaches a plateau at about 8.5. The variabilities of the BCR affinities, as defined by the 25-75% quantiles and 2.5-97.5% percentiles, also shift upward and then plateau along with the mean, despite that the affinities of some B cells reach as low as near 2 and as high as over 12 at times. PCs start to emerge shortly after 20,000 *mcs*, with >10 affinity levels (Fig. 4A, 10 is defined as the threshold affinity to trigger terminal differentiation in the model). The production rate of PCs continues to increase albeit in a highly stochastic fashion and the total cumulative number of PCs produced at the end of GC reaches 2000 (Fig. 4B). Only a tiny fraction of the B cell population becomes PCs in each 600 *mcs* time period, reaching as high as 2% at the end of simulation (Fig. 4C). The PC antibody affinities range from 10 to 12.5, occupying the right tail of the over affinity distribution (Fig. 4D). For those B cells that are positively selected, their mean affinity is 8.28, which is higher than the mean affinity 7.22 of those negatively selected cells. For those B cells that die due to no access to Tfh cells, their affinities cover a broader range on the high end, some of which reach 12.5. Among the initial 200 B cell clones, only 6 clones remain at the end (Fig. 4E), among which one single clone dominates, comprising over 60% of the total B cells at 72,000 *mcs*, followed by two other clones each comprising about 12%, while the remaining 3 clones are much smaller (Fig. 4F). Additional simulations showed that the fractions of dominant clones may vary for each GC – in some cases a single clone absolutely dominates the GC, occupying nearly 90% of the B cells (Fig. S1A and S1B), while in other cases the GC can be co-inhabited by several clones with no single dominant clone (Fig. S1C and S1D).

**Figure 4.**
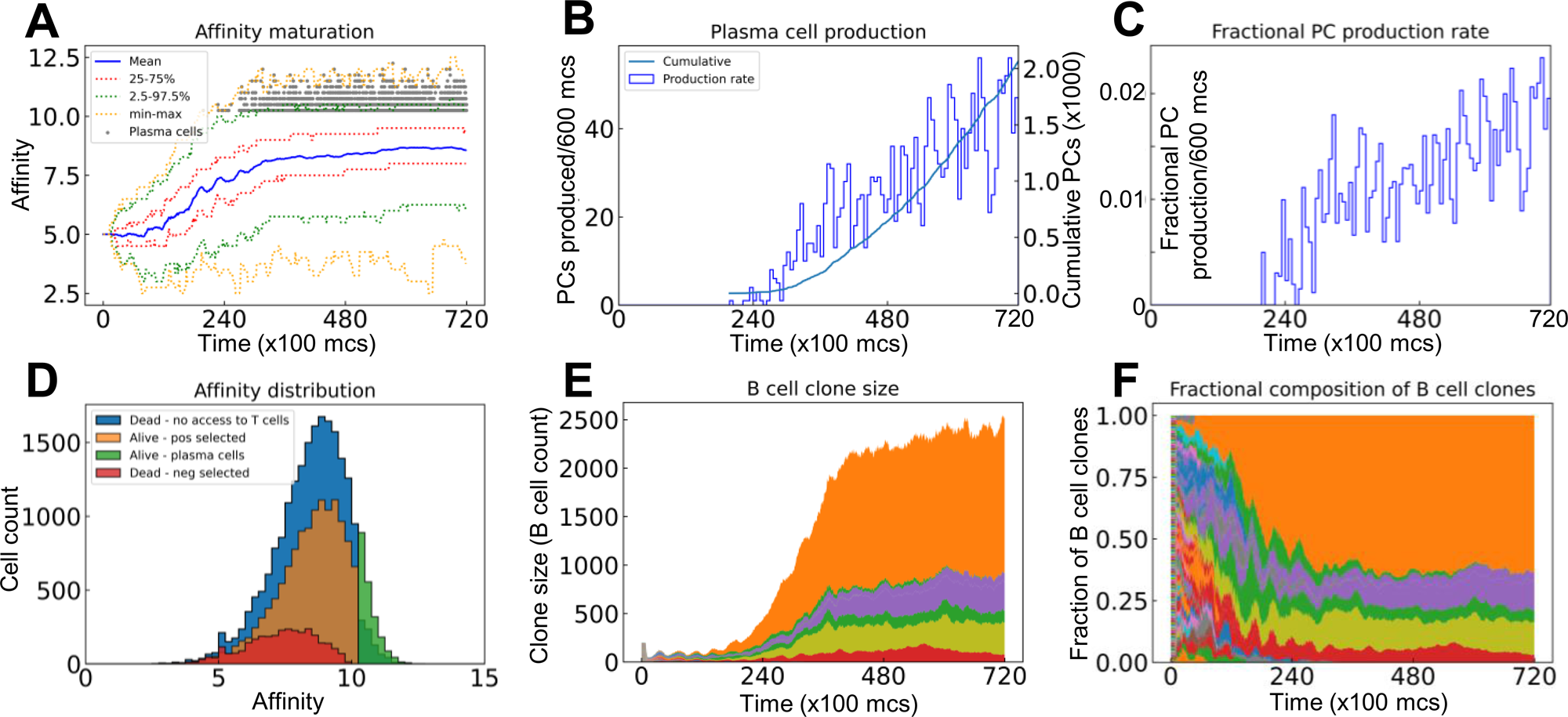
Evolution of B cell affinity maturation and clonal dominance. **(A)** Mean, interquartile, 2.5-97.5^th^ percentile, minimum and maximum BCR affinities of GC B cells as indicated. Gray dots denote the time and antibody affinities of PCs when they emerge. **(B)** Numbers of PCs produced every 600 *mcs* and cumulative numbers of PCs produced as indicated. **(C)** Fractions of B cells that differentiate into PCs in every 600 *mcs*. **(D)** Distributions of BCR/antibody affinities in B cells or PCs as indicated. **(E)** Evolution of B cell clone size as represented by the number of progeny B cells descending from each of the initial 200 clones. **(F)** Muller plot of evolution of the clonal fractions of the GC B cells.

#### 2.3 Inter-zonal migration and LZ residence time

We next characterized the statistics of B cell migrations between the DZ and LZ. For the GC simulation presented in Fig. 2, there are a total of 33,440 B cells that enter from the DZ to the LZ, among which 54.6% die, 37.0% are positively selected and return to the DZ (thus making a full DZ-LZ-DZ round trip), 6.3% differentiate into PCs, and the rest remain in the LZ within the 72,000 *mcs* timeframe of the simulation (Fig. 5A).

**Figure 5.**
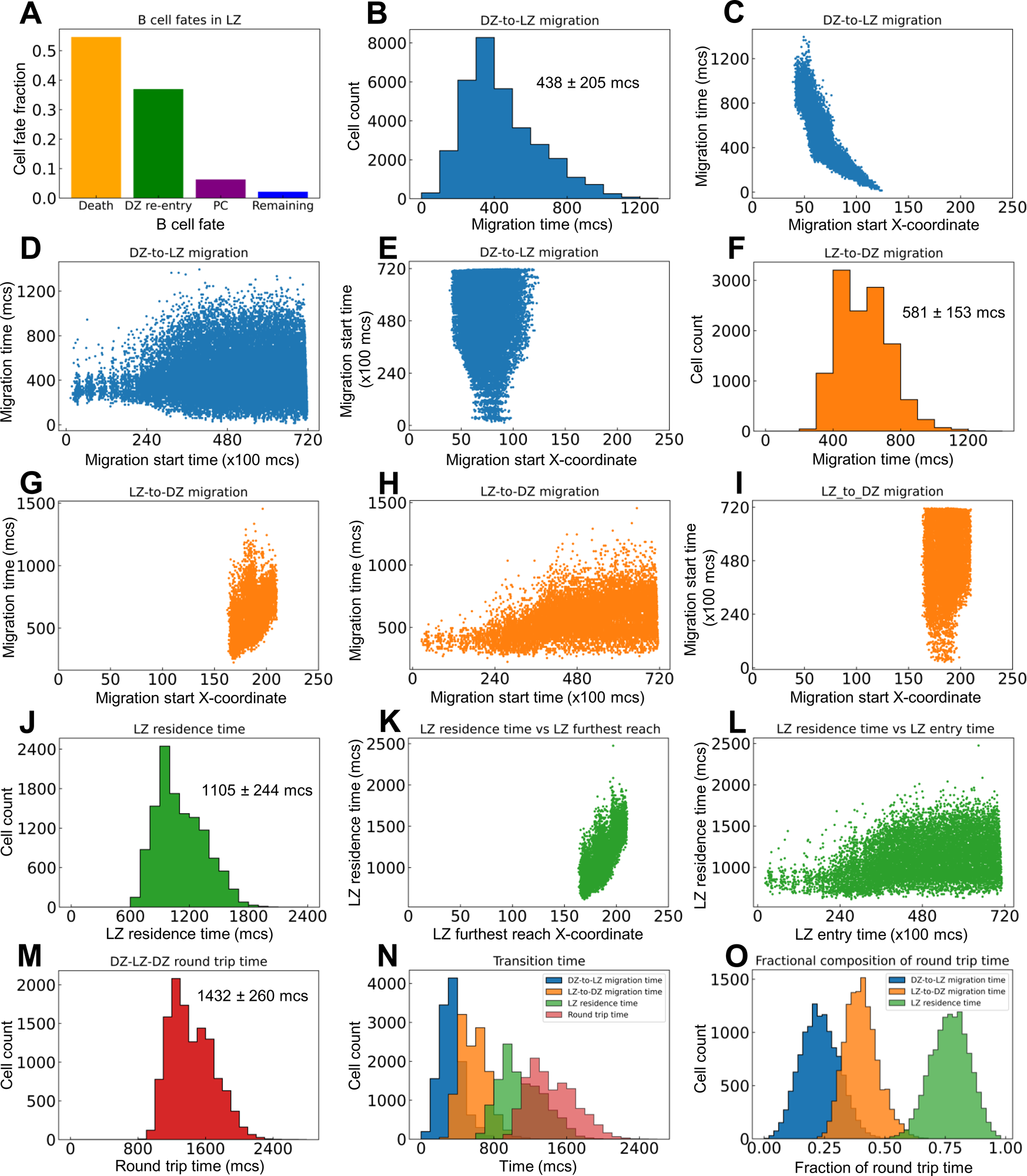
Quantitative analyses of inter-zonal migration. **(A)** Fractions of fates of B cells that have entered the LZ. **(B)** Distribution of DZ-to-LZ migration time (*T_DL_*) with mean ± std indicated. **(C)** Relationship between *T_DL_*and X-coordinate of DZ-to-LZ migration start location. **(D)** Relationship between *T_DL_* and DZ-to-LZ migration start time. **(E)** Relationship between DZ-to-LZ migration start time and location. **(F)** Distribution of LZ-to-DZ migration time (*T_LD_*) with mean ± std indicated. **(G)** Relationship between *T_LD_* and X-coordinate of LZ-to-DZ migration start location. **(H)** Relationship between *T_LD_* and LZ-to-DZ migration start time. **(I)** Relationship between LZ-to-DZ migration start time and location. **(J)** Distribution of LZ residence time (*T_LZ_*) with mean ± std indicated. **(K)** Relationship between *T_LZ_* and X-coordinate of LZ furthest reach of the B cells. **(L)** Relationship between T*_LZ_* and LZ entry time. **(M)** Distribution of DZ-LZ-DZ round trip time (*T_DLD_*) with mean ± std indicated. **(N)** Overlay of distributions of *T_DL_*, *T_LD_*, *T_LZ_*, and *T_DLD_*. **(N)** Distributions of *T_DL_*, *T_LD_*, and *T_LZ_* as fractions of *T_DLD_*.

For the B cells that make a full DZ-LZ-DZ round trip, we calculated the durations they spend in different legs of the trip. The DZ-to-LZ migration time (*T_DL_*) is defined as the duration from the moment a B cell exits the cell cycle in the DZ and starts to migrate to the LZ to the moment the cell crosses the midline, i.e., the DZ/LZ border defined here. *T_DL_* follows a right-skewed distribution, with the median, mean, and standard deviation (std) at 390, 438, and 205 *mcs* respectively (Fig. 5B). *T_DL_*is inversely correlated with the X-coordinate where the DZ-to-LZ migration is started (Fig. 5C), which is expected because migrations initiated further away from the DZ/LZ border will take a much longer time to complete than those initiated near the border. The variability of *T_DL_* increases as DZ-to-LZ migrations start at later times, broadening to both shorter and longer durations (Fig. 5D). This increased variability can be attributed to the more spread-out X-coordinate of the migration start locations as the GC grows in size over time (Fig. 5E).

Similarly, the LZ-to-DZ migration time (*T_LD_*) is defined as the duration from the moment a B cell is positively selected in LZ and starts the DZ re-entry journey to the moment the cell crosses the midline. *T_LD_*also follows a right-skewed distribution, with the median, mean, and std at 570, 581, and 153 *mcs*, respectively (Fig. 5F). *T_LD_* is positively correlated with the X-coordinate where the LZ-to-LD migration is started (Fig. 5G), i.e., the migrations initiated further away from the DZ/LZ border take a longer time to complete than those initiated near the border. When the LZ-to-DZ migrations start at later times, the distribution of *T_LD_* broadens to the long side, without dropping further on the short side (Fig. 5H). The broadening to longer durations can be attributed to the fact that as the GC grows there are more LZ-to-DZ migrations that are initiated at locations in the LZ further away from the midline (Fig. 5I). In comparison, the lower bound to the LZ-to-DZ migration time can be attributed to the locations of the Tfh cells in the LZ, where the ones nearest to the midline are at X-coordinate of about 160 (Fig. 2A), which is 35 pixels (µM) away from the midline.

The LZ residence time (*T_LZ_*), which contains *T_LD_*, is defined as the duration a B cell spends in the confine of the LZ before it reenters the DZ. Like the other two metrics, *T_DL_* and *T_LZ_*, *T_LZ_* also follows a right-skewed distribution, with the median, mean, and std at 1065, 1105, and 244 *mcs* respectively (Fig. 5J). *T_LZ_*is positively correlated with the X-coordinate of the furthest reach of the B cells in LZ (Fig. 5K), and broadens to longer time as the GC progresses (Fig. 5L). There is no correlation between the LZ entry time and furthest reach in LZ (result not shown).

Lastly, we examined the DZ-LZ-DZ round trip time (*T_DLD_*) by summing *T_DL_* and *T_LZ_*. The median, mean, and std of the right-skewed distribution are 1395, 1432, and 260 *mcs* respectively (Fig. 5M). Examining the distributions of the components of *T_DLD_* indicted that on average a B cell spends the least amount of time migrating from DZ to LZ, followed by a 1.3-fold longer time migrating from LZ back to DZ (Fig. 5N). A B cell spends a much longer time in LZ than in transition between the two zones, such that on average *T_LZ_* is about 1.9 and 2.5-fold longer than *T_DL_* and *T_LD_*, respectively. Overall, the round trip breaks down to 23% of the time spent on DZ-to-LZ migration and 77% on LZ residence among which 40% on LZ-to-DZ migration (Fig. 5O).

#### 2.4 Proliferative burst and affinity

B cells positively selected in the LZ initiate their proliferative burst by entering S phase while migrating back toward the DZ (Victora et al. 2010, Gitlin et al. 2014), and complete the first division of the proliferative burst primarily in the DZ. Since *T_LD_* = 581 ± 153 *mcs*, a time delay of 800 *mcs* for cell growth in the first cell cycle was introduced in the model to ensure that cytokinesis does not occur before B cells reach the DZ. Simulations confirmed that this is indeed the case for the first cell cycles of proliferative bursts of nearly all DZ-returning B cells, while all subsequent cycles in the bursts are started and completed within the DZ (Fig. 6A). In a proliferative burst, the length of the first cell cycle is 1412±190 and those of subsequent cycles are 592±82 *mcs* respectively (Fig. 6B). Putting these numbers into perspective, the subsequent cycles have an average length equivalent to nearly 6 hours. The variability of cell cycle length increases as the GC progresses, particularly to the long side (Fig. 6C), suggesting that in a more crowed space, more time is needed to allow “pixel copy” to occur in the CompuCell3D environment in order for cells to grow to the pre-cytokinesis volume. The X-coordinate locations at which cell cycles are initiated expand, especially for subsequent cycles in a proliferative burst (Fig. 6D). The average proliferative burst size, i.e., the number of cell divisions in each burst, is 2.19±0.88 (Fig. 6E). While the majority of B cells that return to the DZ can mount a burst of 2 divisions, some go on to divide up to 5 divisions. Examining the relationship between the affinity of each DZ-returning B cell and its burst size (Fig. 6F) revealed that while a high affinity does not guarantee a large burst size (due to the randomly produced damaging BCR mutations that would kill the daughter cells and thus terminate the burst prematurely), the affinity appears to be positively correlated with the highest achievable burst size. When the affinities are binned into different intervals: <5, 5-6, 6-7, 7-8, 8-9, and >=10, the burst size at 95 percentile in each bin, which are likely the proliferations that contribute to the dominating B cells in the GC eventually, is positively associated with the affinity (Fig. 6G). The mean burst size in each affinity bin also exhibits a positive albeit less strong relationship with the affinity (Fig. 6H). Higher affinities correlate with longer LZ residence time (Fig. 6I and 6J), which is likely a result of association rather than causality, since improvement in affinity and GC population growth occur simultaneously over time, and a denser GC population at a later time tends to result in a longer navigating time through the LZ due to crowding (Fig. 5L).

**Figure 6.**
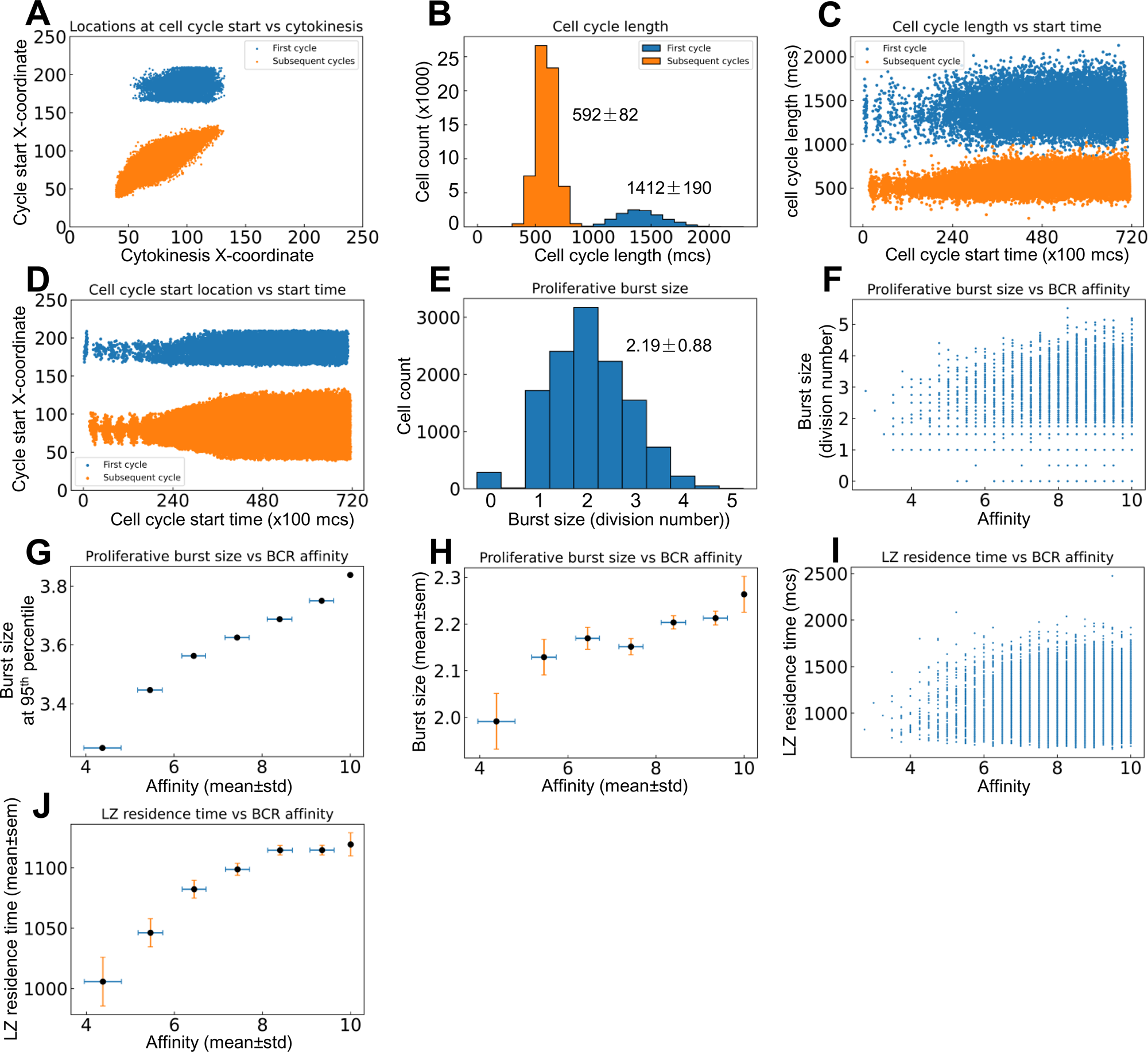
Quantitative analyses of proliferative bursts and BCR affinities of GC B cells. **(A)** Relationship between the X-coordinates of cell cycle start location and cytokinesis location for the first and subsequent cycles as indicated in a proliferative burst. **(B)** Distributions of cell cycle lengths of the first and subsequent cycles in a proliferative burst. **(C)** Relationship between cell cycle length and start time of the first and subsequent cycles in a proliferative burst. **(D)** Relationship between cell cycle start location and start time of the first and subsequent cycles in a proliferative burst. **(E)** Distribution of proliferative burst size. **(F)** Relationship between the proliferative burst size and affinity of DZ-returning B cells. **(G)** Association between the 95^th^ percentile proliferative burst size and mean affinity of DZ-returning B cells in each affinity bin as indicated. **(H)** Association between the mean proliferative burst size and mean affinity of DZ-returning B cells in each affinity bin as indicated. **(I)** Relationship between the LZ residence time and affinity of DZ-returning B cells. **(J)** Association between the mean LZ residence time and mean affinity of DZ-returning B cells in each affinity bin as indicated.

### 3. Molecular responses of B cells and their relationships with cellular phenotypes

The above patterns of phenotypical behaviors of B cells are underpinned by the molecular network operating in each B cell responding to extracellular signaling cues in the GC, including chemoattractants CXCL12 and 13, BCR-mediated antigen signaling by engaging FDCs, and CD40 signaling by engaging Tfh cells (Fig. 1). Examining the steady-state GC population at 700,000 *mcs* revealed that the expression or activity levels of the signaling molecules and genes in the 2352 B cells in the DZ and LZ are highly heterogenous (Fig. 7A). While there is no pMHCII expression in the DZ, a large fraction of B cells in the LZ expresses high pMHCII levels (Fig. 7B), and those expressing low pMHCII levels are B cells that either have just entered the LZ or are en route returning to the DZ with pMHCII being downregulated. A small fraction of B cells in the LZ exhibits high AKT levels as they interact with FDCs (Fig. 7C), which causes downregulation of FOXO1 (Fig. 7D). Signaling downstream of CD40 signaling are cRel and MYC, which are active in a small fraction of B cells in the LZ (Fig. 7E-7G). MYC activity induces AP4, which remains upregulated for an extended period of time, even after the B cells have returned to the DZ and MYC has been downregulated (Fig. 7H), to sustain proliferative bursts. The majority of B cells in the DZ express CXCR4 and the small fraction with CXCR4 downregulated are B cells that have exited the cell cycle and are about to migrate to the LZ (Fig. 7I). There is no difference in CXCR5 expression in B cells between the two zones (Fig. 7A). A few cells in the LZ exhibit high RelA (Fig. 7A) and BLIMP1 (Fig. 7J) expression representing emerging PCs. The correlations between select pairs of the signaling molecules are shown in Fig. 8. There is a strong negative correlation between FOXO1 and AKT (Fig. 8A), positive correlations between cRel and CD40 (Fig. 8C), MYC and CD40 (Fig. 8D), MYC and cRel (Fig. 8E), and AP4 and MYC (Fig. 8F) in B cells in the LZ, while CXCR4 is only expressed in high FOXO1-expressing B cells (Fig. 8B).

**Figure 7.**
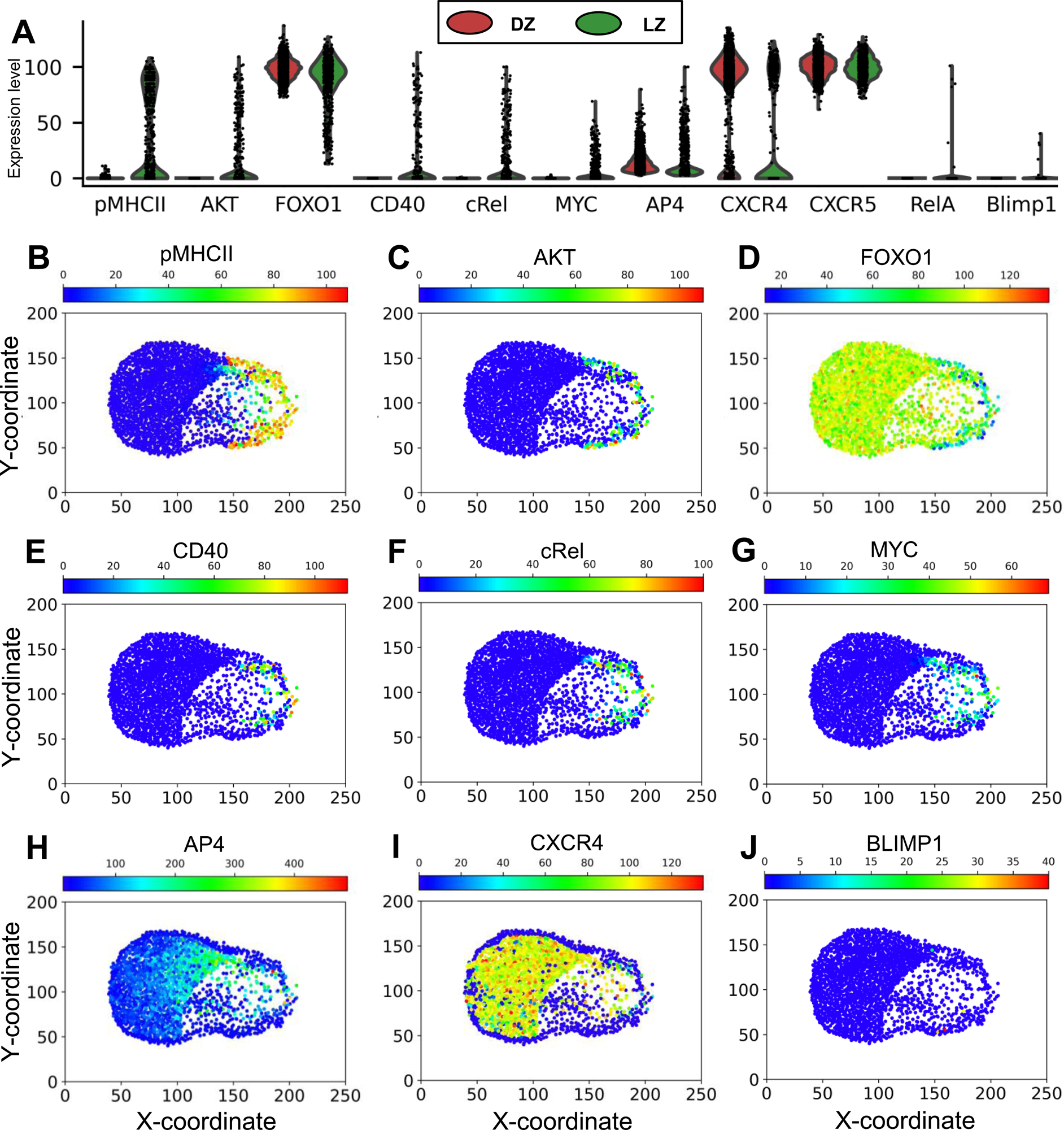
Molecular response profiles of GC B cells. **(A)** Violin plots of expression/activity levels of signaling molecules in DZ and LZ B cells as indicated at 70,000 *mcs*. **(B)** Simulated “immunohistochemistry” staining of signaling molecules as indicated in GC B cells at 70,000 *mcs*.

**Figure 8.**
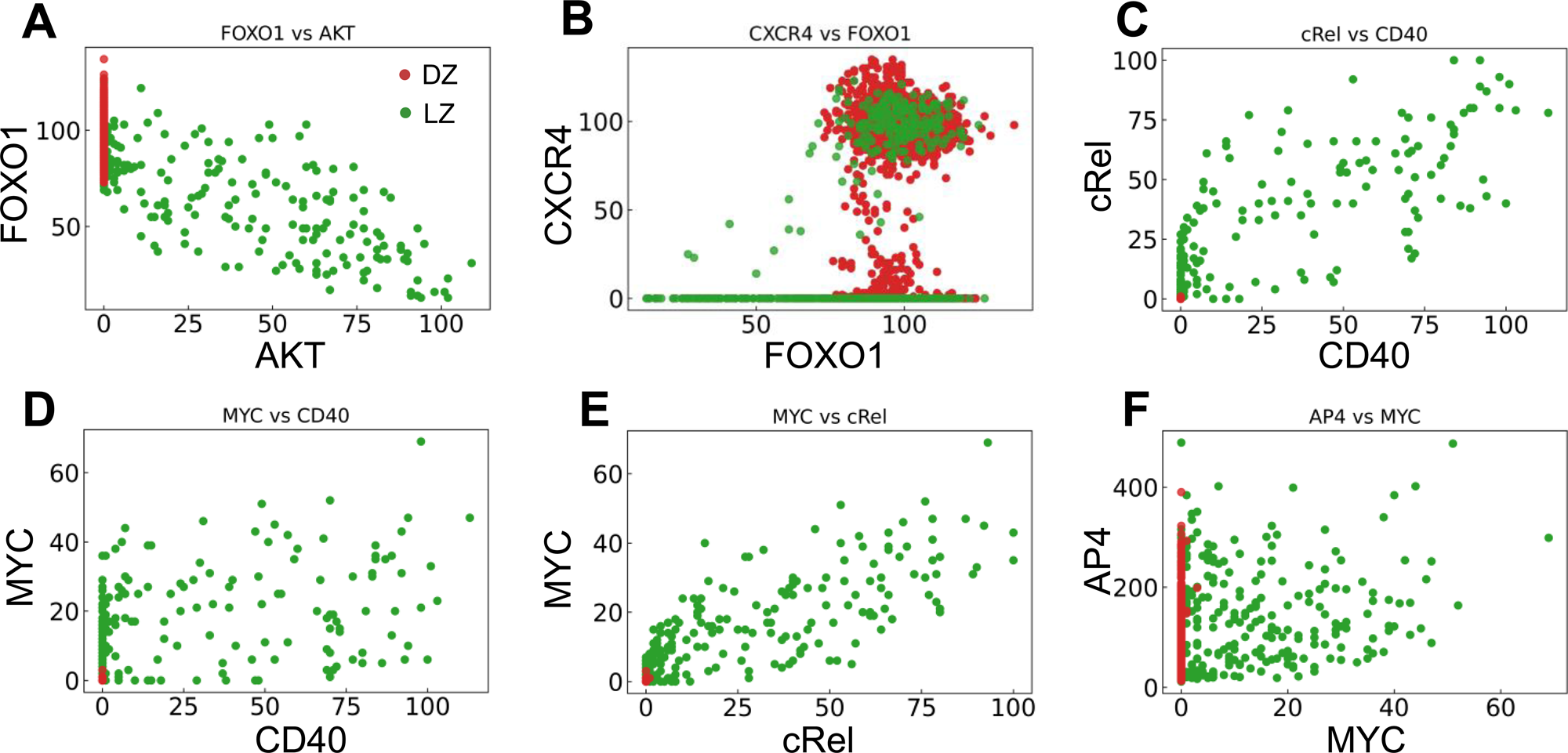
Correlations between key signaling molecules in DZ and LZ B cells as indicated at 70,000 *mcs*. Red and green dots denote DZ and LZ B cells respectively as shown in (A).

We next examined the relationship between the signaling molecules and phenotypical behaviors of B cells that are positively selected in the LZ and return to the DZ. Both the peak level (Fig. 9A) and duration of MYC expression (Fig. 9B), which is quantified by the area under the curve (AUC), are positively correlated with the BCR affinity of the cells. The correlation indicates that the strength of the affinity-dependent BCR signaling, which downregulates FOXO1 and upregulates pMHCII to enable Tfh-dependent CD40 signaling, is quantitatively transmitted to MYC. Although MYC is only transiently expressed in the positively selected B cells in the LZ, its encoding of affinity is relayed to AP4. By integrating the MYC signal, AP4, which has a longer half-life than MYC, can be activated for a much longer time as B cells migrate back to the DZ. Its peak level is positively correlated with the AUC of MYC in DZ-returning B cells (Fig. 9C). The 95^th^ percentile burst size is positively correlated with the mean AP4 peak level in each affinity bin (Fig. 9D).

**Figure 9.**
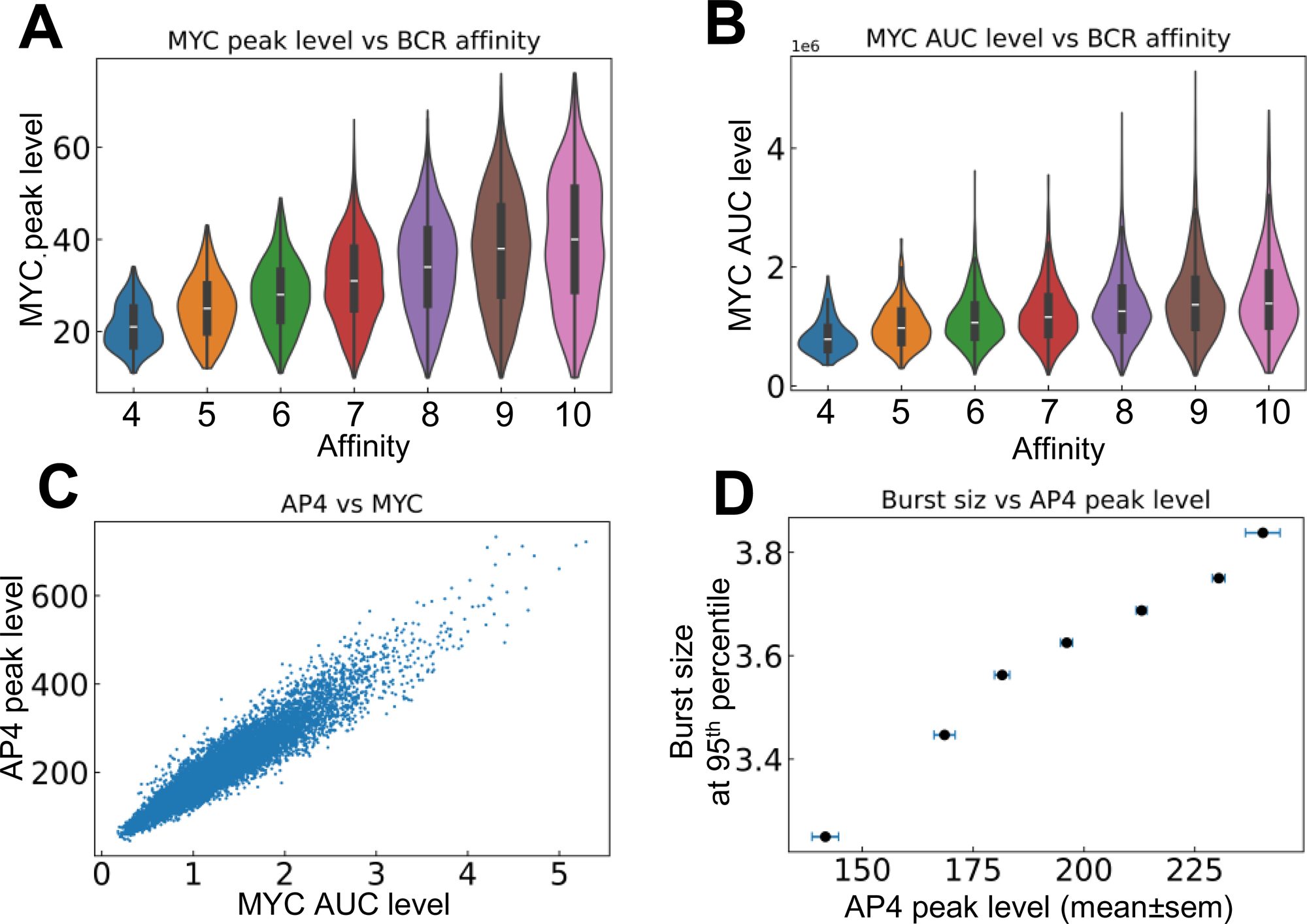
Relationships between signaling molecules and phenotypical behaviors of DZ-returning B cells. **(A)** Violin plots of MYC peak levels in different affinity bins. **(B)** Violin plots of MYC AUC level in different affinity bins. **(C)** Correlation between AP4 peak level and MYC AUC level. **(D)** Correlation between the 95^th^ percentile proliferative burst size and mean AP4 peak level in each affinity bin.

Lastly, the trajectory of a representative single B cell lineage branch that successfully makes it to a PC is presented (Fig. 10). As the seeding B cell and its progeny cycle between the DZ and LZ (Fig. 10A), multiple cell divisions (between 3-6 divisions) occur in each proliferative burst (Fig. 10B) with increasing cell generation (Fig. 10C). In this particular simulation result, nearly every cell division results in an increase in BCR affinity despite a few occasions when the affinity decreases (Fig. 10D). Each time the B cell starts the trip to the LZ the countdown of the death timer is initiated, but in this case it never drops below the predefined death threshold of 50 before the cell is rescued by positive selection where the death timer is reset (Fig. 10E). Every time the B cell moves into the LZ, pMHCII is re-expressed proportionally to the antigen-specific affinity after encounter with FDCs that triggers BCR signaling (Fig. 10F). BCR signaling transiently activates AKT (Fig. 10G) which downregulates FOXO1 transiently (Fig. 10H). Commitment to cell cycle after the B cell is positively selected allows upregulation of CXCR4 by FOXO1, which drives the B cell to return to the DZ, and CXCR4 is downregulated after the B cell exits the last cell cycle of a proliferative burst (Fig. 10I), which allows the cell to migrate to the LZ due to constitutively expressed CXCR5 (not shown). In the LZ, encounter with Tfh cells triggers transient activation of the CD40-cRel-MYC axis in the presence of downregulation of FOXO1 (Fig. 10J-10L). By integrating the MYC signal, AP4 is upregulated for an extended period of time, which lasts well into the DZ (Fig. 10M) to sustain the proliferative bursts. The proliferative bursts terminate when AP4 drops below a predefined threshold of 50. When the affinity increases past a predefined threshold of 10, the BCR signaling triggers RelA activation in a switch-like manner (Fig. 10N), which in turn activates BLIMP1 that drives the B cell to terminally differentiate into a PC (Fig. 10O). A representative result of a B cell lineage that ends in death in the DZ due to damaging BCR mutations is shown in Fig. S2A-S2C, where at the time the damaging mutation occurs the affinity drops to “-1” as an indication, and caspase 3 is upregulated to trigger apoptosis. A representative result of a B cell lineage that ends in death in the LZ because of not being positively selected is shown in Fig. S2D-S2F, where the death timer dips below the threshold of 50 thus triggering cell death.

**Figure 10.**
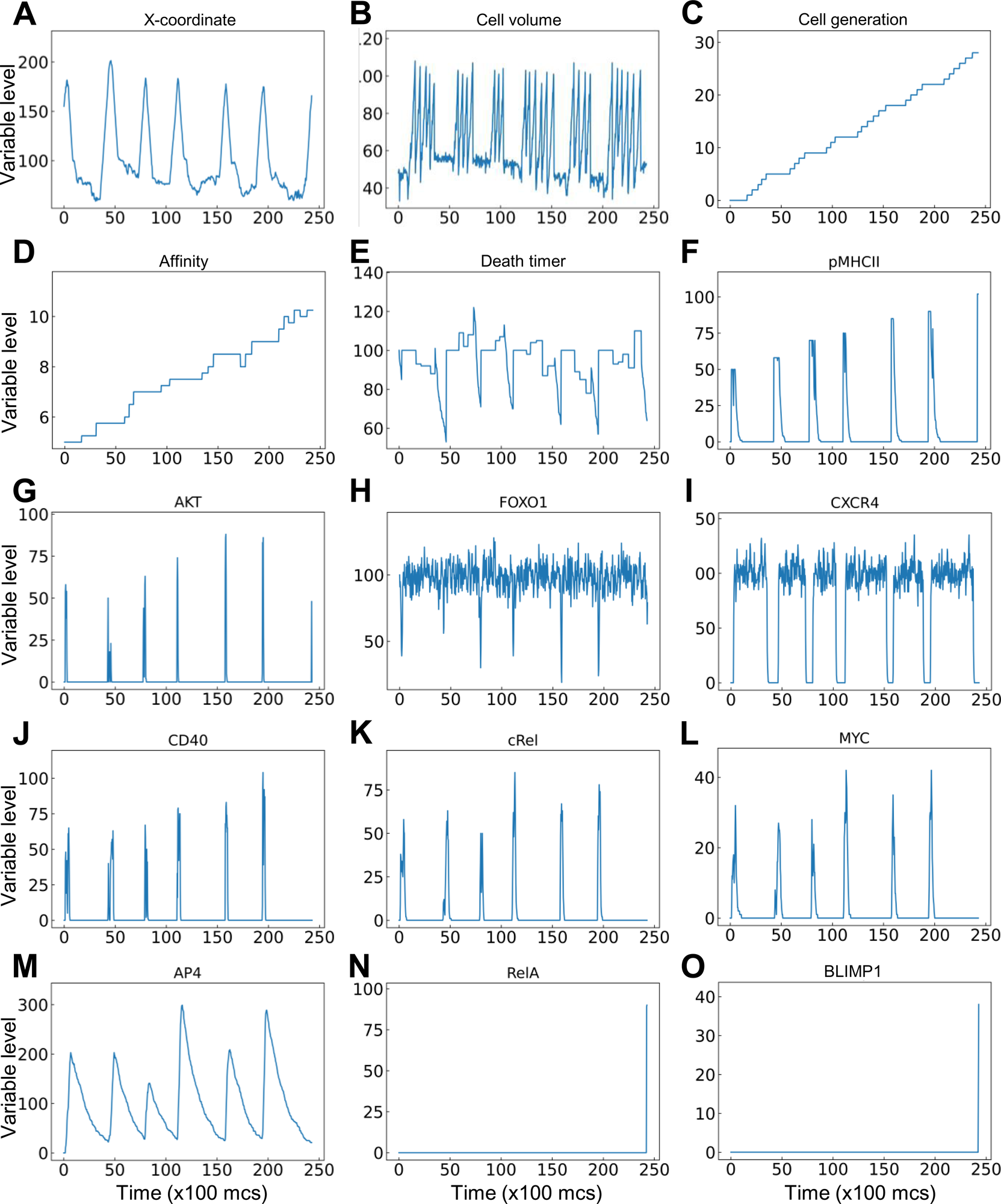
Trajectory of a single B cell lineage branch that successfully makes it to a PC with molecular and cellular variables as indicated.

## Discussion

In the present study, we developed a multiscale spatial modeling framework for the GC in the CompuCell3D simulation platform. By integrating interactions across molecular, cellular, and tissue scales, the model captures key hallmark GC events, including cyclic migration of B cells between the DZ and LZ, proliferative burst, SHM, deaths due to damaging BCR mutation, positive selection, timed cell death, affinity maturation, clonal expansion and dominance, and loss of clonal diversity. These cellular behaviors are driven by simulating an underlying molecular network in individual B cells responding to BCR and CD40 activation via interacting with FDCs and Tfh cells, respectively. The molecular outputs of the network include CXCR4 driving LZ-to-DZ chemotaxis, MYC and AP4 driving cell cycles, and caspase 3 driving apoptosis. While this multiscale model can be further tuned and elaborated to study diverse variables and conditions regulating GC outcomes, exploring these possibilities is beyond the scope of the present study. We focused on demonstrating the capability of the modeling framework by presenting a simulated GC that leads to affinity maturation, with qualitative and to some extent quantitative results that are commensurate with the primary literature.

### 1. GC B cell birth, death, and population dynamics

Emerging GCs are seeded with a few hundred B cell clones (Wittenbrink et al. 2010). Starting with 200 B cells, our GC simulation showed that the DZ:LZ ratio of the numbers of B cells oscillates at early time points (Fig. 3A). This occurs because there are not many B cells at this stage of the GC and these cells tend to migrate in sync between the two zones. As the GC B cell population approaches a steady state, the DZ:LZ ratio converges to a constant value, nearly 2.6:1, in our simulation, which is comparable to the 2.15∼2.2:1 ratio observed in both mouse and human GCs (Victora et al. 2012). The total number of B cells in the simulation can reach about 2500. Given that the modeled GC space contains only 11 pixels (equivalent to 11 µM) along the Z-dimension to keep the simulation time tractable, we expect that when scaling up by 2-5 times to mimic the actual GC thickness of 20-50 µm (Olivieri et al. 2013), the peak number of B cells will reach thousands to over 10,000, consistent with the estimated B cell counts in real GCs (Wittenbrink et al. 2010).

Our simulation showed that new B cells are born in the DZ, while cell deaths occur in both the DZ and LZ at an overall rate that eventually matches the birth rate as the GC approaches the steady state. Although the absolute number of deaths in the DZ is more than twice that in the LZ (Fig. 3I), the percentage death rates are comparable, at about 40-43% per 600 *mcs* in both zones (Fig. 3J). These numbers are concordant with the nearly 50% death rate per 6 hours reported for the GCs in Peyer’s patches in mice and GCs in mice immunized with 4-hydroxy-3-nitrophenylacetyl (NP)-conjugated ovalbumin (NP-OVA) or HIV-1 envelope antigen GT1.1 (Mayer et al. 2017). The steady-state death turnover rate of 50% per 6 hours is expected given that GC B cells proliferate with a cell-cycle length of 4-6 hours (Mintz and Cyster 2020, Victora and Nussenzweig 2022) and in our model the cell volume doubling time of proliferating B cells is parameterized at 500 *mcs*. The apoptotic deaths in the DZ are believed to occur in the late G1 phase, triggered by AID-induced BCR-damaging mutations during transcription, including stop codons, insertions and deletions in the Ig sequences, such that the synthesized BCR proteins fail to properly fold and be expressed on the cell surface (Mayer et al. 2017, Stewart et al. 2018).

In the LZ, the default fate of B cells is apoptosis if not positively selected regardless of their BCR affinity, and the apoptosis will occur when the death timer goes off, which is initiated after the B cells exit the cell cycle in the DZ (Heinzel et al. 2017, Mayer et al. 2017). Our simulations showed that only a small fraction of deaths in the LZ results from not being positively selected after contact with FDCs and Tfh cells, while the majority of the deaths are due to no access to Tfh at all (Fig. 3M). This result recapitulates the current notion that the availability of Tfh cells is the limiting factor, not the competition for antigen, for positive selection (Meyer-Hermann et al. 2006, Allen et al. 2007b, Meyer-Hermann 2007, Victora et al. 2010). Interestingly, our simulations also suggested that at the advanced GC stage, a small fraction of cell deaths in the DZ may also result from the lack of access to Tfh if these cells do not have enough time to migrate through an increasingly densely populated DZ to reach the LZ (Fig. 3N and 3O). Whether deaths of such nature occur *in vivo* and to what extent remain to be tested experimentally.

The dynamics of the GC B cell population is determined primarily by the cell birth rate and death rate. The loss of B cells as a result of GC exit as memory and PCs is expected to be negligible because they only account for <3% of the GC B cell fates (Kräutler et al. 2017, Laidlaw et al. 2017, Holmes et al. 2020). Our simulation result is consistent with this estimate: PCs emerge at a fraction of only as high as 2% of the B cell population towards the end of the simulated GC (Fig. 4C). For a GC B cell population to grow or to avoid population collapse, the average proliferation rate has to be higher than the death rate. Absent any limiting factors, the cell population dynamics in the GC may operate as a positive feedback system, producing bistability of two alternative outcomes – expansion or regression – as predicted by previous mathematical models (Meyer-Hermann and Beyer 2004). In the expansion mode, as the overall affinities increase, the probability of cell death in the LZ owing to lack of positive selection decreases, thus more B cells will return to the DZ and proliferate with a larger burst size there, which results in a higher birth rate of progeny cells with potentially even higher affinities returning to the LZ, and the cycle repeats leading to GC B cell expansion. In the regression mode, the positive feedback works in the opposite direction, where lower affinities can lead to fewer B cells positively selected in the LZ and smaller burst size in the DZ, which eventually leads to GC regression. This all-or-none type of GC outcomes is consistent with the population bottleneck proposition (Zhang and Shakhnovich 2010) and suggests that there could be an initial affinity threshold condition for those activated B cells that seed a GC, below which a tangible GC is unlikely to emerge and above which a GC will likely emerge and grow. This suggests that it may take B cells of some intermediate affinity to initiate a GC to produce optimal humoral immune outcomes, i.e., high production of high-affinity PCs. A GC starting with B cells of too low affinity will likely abort prematurely as argued above, while a GC starting with high-affinity B cells will likely grow but only to a small size before some B cells hit the affinity threshold that triggers terminal differentiation to PCs. This may also explain the observation that affinity selection for memory B cells is less stringent, and they are often formed and then exit the GC when their affinities are still low or intermediate levels, while the PCs are in general of high affinity (Smith et al. 1997, Phan et al. 2006, Shinnakasu et al. 2016, Kräutler et al. 2017, Viant et al. 2020). Upon secondary infection or booster immunization whether the recalled memory B cells directly differentiate to PCs or have to go through GCR again can be determined by many factors (Inoue et al. 2018, Valeri et al. 2022), and it is likely that their BCR affinity may play a role in this regard.

When a GC reaches a certain size, its growth could be restricted by multiple factors, before other GC-shutoff mechanisms such as antigen depletion and antibody feedback kick in. In the present study, we showed that the availability of Tfh cells is a limiting factor, where a higher fraction of B cells in the LZ die because of a lack of access to Tfh cells, thus increasing the overall death probability which balances out the increasing birth rate due to improved antigen affinity. That B cells become PCs once their affinities reach a threshold also helps the GC B cell population to reach an equilibrium by curbing further increase in the number of higher-affinity B cells and thus attenuating the positive feedback mechanism described above. Another limiting factor not considered in the present study is the limited nutritional and energetic resources that could restrict B cell proliferation once the GC has grown to a mature size (Wittenbrink et al. 2010).

To grow a GC the overall damaging BCR mutation-induced cell death probability cannot be higher than 50% for each cell division. Actually it has to be much lower than 50% because not all but only a small fraction of B cells arriving in the LZ are positively selected and return to DZ. In our model, we encoded a DZ death probability of 30% for each of the two daughter cells born from a cell division, which leads to a probability of 49% (0.7*0.7) to double the number of B cells after each division, of 42% (2*0.3*0.7) to keep the number of B cells constant, and of 9% (0.3*0.3) to eliminate the proliferating B cell. The DZ re-entry probability was estimated to be between 10-30% through mathematical modeling and analysis of experimental data (Victora et al. 2010, Meyer-Hermann et al. 2012, Mesin et al. 2016). These values suggest that absent any DZ death, the average proliferative burst size has to be greater than 1.73-3.32 divisions to grow a GC. When taking into consideration DZ death, which is very significant, on par with LZ death (Mayer et al. 2017), the average burst size has to be much higher. The DZ re-entry probability depends on the BCR antigen affinity, thus it is likely that at the early stage of GC the re-entry probability is low and at the advanced stage it is high. In our model, the positive selection and thus DZ re-entry probability is set to be proportional to pMHCII, such that when the affinity is intermediate at 5, the probability is 50% and when the affinity approaches 10 or higher, the DZ re-entry probability is 100%. However, the overall DZ re-entry fraction is only 36% in our simulation (Fig. 5A), which can be attributed in part to the inaccessibility to Tfh cells. A re-entry fraction of 36% requires at least a burst size of 1.47 to grow the GC in the absence of DZ death. With a DZ death probability of 30% for each new born B cell, our model has an average burst size 2.2 (Fig. 6E), which is in general agreement with the estimated average of 2 divisions (Meyer-Hermann et al. 2012, Gitlin et al. 2014, Meyer-Hermann 2021) or 3 divisions per burst (Gitlin et al. 2015) in the literature.

### 2. Affinity maturation, proliferative burst, clonal expansion and dominance

While the average proliferative burst size is 2-3, each burst can vary between 1-6 divisions (Meyer-Hermann et al. 2012, Gitlin et al. 2014, Gitlin et al. 2015, Meyer-Hermann 2021). This range is quantitively captured in the burst size distribution produced by our simulation (Fig. 6E). The right-tailed burst size distribution could result intrinsically and in part from the probabilistic damaging mutation-induced cell death after each cell division, which increases the chance of short bursts but limits the highest attainable number of divisions in a proliferative burst even for high-affinity B cells. GC B cells positively selected are guaranteed to divide once, while the number of additional divisions or the burst size is directly proportional to the amount of antigen captured by B cells from FDCs and presented to Tfh cells (Gitlin et al. 2014, Finkin et al. 2019). Our model reproduces this positive association (Fig. 6G-6H). Moreover, because of potential premature termination of a proliferative burst induced by damaging mutation, the affinity appears to be better correlated with the top attainable burst size than the average burst size.

The translation of BCR antigen affinity into burst size is mediated molecularly by two key transcription factors: MYC and one of its target genes AP4. Because of the transient nature of B cell interactions with FDCs and Tfh cells (Allen et al. 2007b) and the short half-life of MYC (Heinzel et al. 2017), MYC is only transiently expressed in a small fraction of LZ B cells (Calado et al. 2012, Dominguez-Sola et al. 2012). The MYC expression level is in direct proportion to the amount of antigen captured and dictates the proliferative burst size (Finkin et al. 2019). While MYC can initiate cell growth and cell cycle by driving LZ B cells into the S phase, its lasting effect on cell proliferation is mediated by AP4, which is induced by MYC in a delayed fashion in positively selected GC B cells and is sustained after the B cells re-enter the DZ (Chou et al. 2016). Our model recapitulates the spatiotemporal dynamics of MYC by showing transient MYC expression in LZ B cells, its positive correlation with BCR antigen affinity, and sustained AP4 expression.

Our model recapitulates a typical affinity maturation process along with GC growth (Fig. 4A). The progressive increase in the mean affinity of the GC B cell population is not because all or the majority of the seeding B cell clones improve their affinities uniformly. Rather, in most cases, the mature GC B cell population is dominated by progenies of one or a few of the initial 200 seeding clones (Fig. 4F, S1B, and S1D). This simulation result of terminal clonal dominance is consistent with the premise that GCs mature oligo-clonally (Kroese et al. 1987, Küppers et al. 1993). More recently, using multiphoton microscopy and sequencing Tas and colleagues further revealed that a GC can start with tens to hundreds of distinct B cell clones but loses the clonal diversity over time, converging to one or a few parallelly expanding clones (Tas et al. 2016). The single dominant clone can constitute 10-100% of the final GC B cell population. They further showed that clonal dominance can be achieved through neutral competition, due to stochastic effect, even when all seeding B cells have equal affinity and cannot undergo SHM, a finding that can be explored with our model in the future.

### 3. Inter-zonal migration

Beltman et al. analyzed time-lapsing imaging data of GCs and revealed that B cells move at a net speed of 0.2-0.3 µm/min toward the LZ to produce a DZ-to-LZ migration time of a few hours (Beltman et al. 2011). In our simulation the DZ-to-LZ migration time is about 438 ± 205 *mcs* (Fig. 5B), consistent with that estimated by Beltman. The LZ-to-DZ migration time in our simulation is 581± 205 *mcs* (Fig. 5F) which is about 33% longer than the DZ-to-LZ migration time. The longer time is consistent with the experimental observations (Victora et al. 2010), and could be attributed to the fact that B cells returning to the DZ have to move against the much heavier incoming traffic of B cells migrating from DZ to LZ. The overall LZ residence time is 1105 ± 244 *mcs* (Fig. 5J). These travel times can be tuned by varying the positions and distributions of CRCs and FDCs/Tfh cells in the DZ and LZ respectively, the CXCL12 and 13 gradients, and the chemotaxis strength parameters in the model.

### 4. Existing computational GC models and improvements by our modeling framework

Many mathematical GC models have been developed in the past two decades using a variety of computational approaches, including deterministic, stochastic, agent-based, and hybrid ones implemented in programming languages such as C++, C, MATLAB, and R. Focusing on various aspects of the GCR and simulating at various biological scales, these models have aided our understanding of this long known phenomenon, explored possible modes of mechanisms, predicted GC-associated disease outcomes, and explored optimal design of vaccination schemes. A prominent one among these efforts is the agent-based modeling framework *hyphasma* pioneered by Meyer-Hermann and colleagues that is implemented in C++ (Meyer-Hermann et al. 2009, Meyer-Hermann et al. 2012, Robert et al. 2017). The base model uses stochastic agent-based approach to simulate the movement of each cell as diffusion on a 3D equidistant lattice. Simulating the GC at the cell and tissue levels, the base model and subsequent iterations have been used to explore a range of mechanistic questions, generating novel and useful insights some of which have been validated experimentally.

GC Issues explored with earlier versions of these models include: signaling vs. chemotaxis modes of action (Beyer et al. 2002, Meyer-Hermann 2002, Meyer-Hermann and Beyer 2002), requirement of DZ re-entry for GC development and affinity maturation (Meyer-Hermann and Maini 2005a), whether affinity maturation is driven by competition for antigen or Tfh cells, GC termination resulting from antigen depletion (Meyer-Hermann et al. 2006, Meyer-Hermann 2007), persistent random walk of B cells observed with two-photon imaging and the requirement of active chemotaxis for maintenance of GC zonation (Meyer-Hermann and Maini 2005b, Figge et al. 2008, Binder and Meyer-Hermann 2016), and affinity-dependent proliferative burst size enabling large, high-affinity GC B cell populations (Meyer-Hermann 2014), Issues explored with later iterations of these models include: model variants informed by FOXO1, MYC, and mTOR signaling dynamics on clonal dominance and independent control of B cell selection and division fate decisions (Meyer-Hermann 2021), the effects of periodic cycling of antigen immune complex in FDCs on GC development (Arulraj et al. 2021b), differences in the lifetime of individual GCs resulting from variations in antigen availability and founder cell composition (Arulraj et al. 2021a), contribution of GC-GC interactions to variability in the timing of individual GC maturation (Arulraj et al. 2022a), different mechanisms of GC shutdown (Arulraj et al. 2022b), the effects of kinetic rates of BCR-antigen binding on antigen uptake by B cells and GC dynamics and outputs (Lashgari et al. 2022), and the evolution of clonal diversity and dominance and the modulating effects of antigen amount, Tfh cell availability, and seeding B cell affinity (Meyer-Hermann et al. 2018, Garg et al. 2023),

Besides the seminal work by Meyer-Hermann and colleagues, many others also modeled the GC with various simulation approaches including deterministic, stochastic, and agent-based ones. With deterministic models, Zhang and Shakhnovich explored the parameter space of mutation rate, selection strength, and initial antigen affinity for maximizing affinity maturation (Zhang and Shakhnovich 2010), Chan et al. showed that the feedback from receptor downregulation induced by the CXCL12 and 13 fields may explain the spontaneous interzonal and intrazonal oscillations of B cells (Chan et al. 2013), and Reshetova et al. revealed there is a limited correlation between the size and antigen affinity of GC B cell subclones and B cells with highest affinity can reside in low-abundance subclones (Reshetova et al. 2017). Beltman et al. constructed a stochastic model that recapitulated the persistent random walk of B cell movement in the GC and the small preference for DZ-to-LZ migration (Beltman et al. 2011). Molari et al. developed stochastic and deterministic models to show that the average GC B cell affinity is determined non-monotonically by the antigen dosage, and clonal dominance and limited diversity can be achieved over time (Molari et al. 2020). Using stochastic models and analytical solutions Molari et al. studied the probability for a B cell lineage to surpass the population bottleneck as a function of the antigen concentration and initial B cell population size (Molari et al. 2021). More recently Yan et al. developed a spatiotemporal stochastic model to understand the determinants of GC size and found there is a critical GC volume to achieve best performance (Yan et al. 2022). With an agent-based modeling framework of Basic Tonsil Unit, Hawkins et al. showed that the persistent random walk of B cells could be an emergent outcome of mobile but morphologically rigid B cells in a GC of dense cellularity, where cells are constantly competing for space (Hawkins et al. 2011). Using a spatiotemporally resolved stochastic model similar to the agent-based model by Meyer-Herman et al, Wang et al. showed the importance of efficient Tfh cell delivery for affinity maturation, suggesting that antagonism between BCR signaling and Tfh cells may accelerate affinity maturation (Wang et al. 2016). Using an agent-based model, Amitai explored the GC population dynamics and diversity of clonal dominance with either birth- or death-limited selection (Amitai et al. 2017).

In addition to probing basic GC biology, computational GC models have also been used to help optimize the design of vaccination schemes. Using a stochastic agent-based model, Wang et al. predicted that for GCs to produce cross-reactive antibodies against different antigen variants, sequential rather than simultaneous vaccination with several antigen variants is preferred, and the *in silico* prediction was validated in mice vaccinated with variant gp120 constructs of the HIV envelope protein (Wang et al. 2015, Wang 2017). Meyer-Hermann showed that a feedback imposed by preexisting antibodies or memory B cells can mask the immunodominant epitopes to diversify GCs toward less frequent epitopes to help generate broadly neutralizing antibodies (Meyer-Hermann 2019). Garg et al. developed a stochastic GC model to explain and predict that passive immunization can promote and optimize GCR by tuning the administered external antibodies to control antigen availability such that only high-affinity B cells prevail (Garg et al. 2019). With an agent-based GC model, Yang et al. interpreted why 3 doses of mRNA vaccine against the original SARS-CoV-2 strain are required to develop anti-Omicron neutralizing antibodies, which involves enhanced antigen availability and immunodominant epitope masking after the 2^nd^ dose, and expansion of memory B cells targeting subdominant epitopes by the 3^rd^ dose (Yang et al. 2023). Adapting the same agent-based model, Bhagchandani et al. explained why a particular two-shot extended-prime regiment of immunization against HIV is effective in producing high-titer antibodies; the model predicted that it was because the antigen delivered in the second dose can be captured more efficiently as immune complexes, which was verified by experiments (Bhagchandani et al. 2023).

While these models above investigated the complex processes of affinity maturation and B cell population dynamics, no molecular networks were included in them to drive the B cell behaviors and GC evolution. Quantitative multiscale mathematical models of GC dynamics have been proposed as predictive frameworks to translate basic immunological knowledge to practical challenges (Verstegen et al. 2021, Vaidehi Narayanan and Hoffmann 2022). In more recent years, modeling efforts in this direction have emerged. Merino Tejero and colleagues have developed multiscale GC models integrating molecular and cellular responses (Merino Tejero et al. 2021a, Merino Tejero et al. 2021b). Implemented in C++ language, they combined the agent-based model developed by (Meyer-Hermann et al. 2012) and ODE-based gene regulatory network model comprising BCL6, IRF4, and BLIMP1 developed in (Martínez et al. 2012). They used the model to study the role of affinity-based CD40 signaling and asymmetric B cell division in temporal switch from memory B cell to PC differentiation and DZ-to-LZ ratio. Lately they adapted the model to examine the oncogenic effects of genetic alteration of the above key transcription factors on GC-originated diffuse large B cell lymphoma (Merino Tejero et al. 2022). More recently, the model was used to explore the relationship between clonal abundance and affinity as well as affinity variability within B cells from the same clone, with an attempt to make sense of repertoire sequencing data (García-Valiente et al. 2023).

In comparison, the multiscale spatial GC modeling framework we developed here in the CompuCell3D platform further integrates across the molecular, cellular, and tissue scales and offers several improvements. The framework allows the molecular network to drive multiple cellular behaviors, including B cell growth, division, chemotaxis, survival/death, and PC differentiation, which in turn collectively drive GC tissue pattern formation; reciprocally the cell-to-cell interactions between B cells and FDCs and Tfh cells drive the responses of the molecular signaling network in B cells. Novel cross-scale strengths include cell cycle and FOXO1-dependent CXCR4 expression driving DZ-reentry chemotaxis, MYC and AP4-dependent cell growth and division burst, and RelA and BLIMP1-dependent PC differentiation. As more molecular species are added to the network, additional cross-scale integrations will become available. Because the simulated B cells comprise multiple pixels, the model allows recapitulation of B cell morphology during chemotaxis and volume growth during cell cycle as well as better mimicking of cell-cell interaction. For future iterations of the model, the CompuCell3D platform can easily include paracrine signaling by ILs and other cytokines secreted by B cells, Tfh cells, and FDCs. Last but not least, with the modular plugins and systems biology markup language (SBML) support, the CompuCell3D platform allows a more structured construction of the GC model that will facilitate future model sharing and integration.

### 5. Limitations and future iterations

As an initial effort to establish the multiscale GC modeling framework on the CompuCel3D platform, we simulated the GC to the mature stage where the B cell population approaches a steady state. While persistent GCs can exist for months or years under chronic infections such as certain viral infections (Bachmann et al. 1996, Kasturi et al. 2011, Adachi et al. 2015), and in the Peyer’s patch for mucosal immunity (Reboldi and Cyster 2016), in most other infection or immunization scenarios, GCs eventually regress in 3-4 weeks with mechanisms not well understood. Some GC terminations may be because the antigens stored in FDCs are depleted, changes in the signaling nature of Tfh cells and FDCs, or antibodies produced by the departed PCs circulate back into the GC and block antigen presentation (Zhang et al. 2013, Arulraj et al. 2021c, Arulraj et al. 2022b). In addition, in the current model, memory B cells are not included as a cell fate option. Since like PCs, memory B cells only constitute a very small fraction of the GC B cells fates (Arulraj et al. 2021c), its exclusion is not expected to affect the overall model behavior. Nonetheless, future iterations of the GC model will include the self-termination and memory B cell formation as driven by the IRF4-BCL6-BLIMP1 network (Martínez et al. 2012), and if needed, CSR, which is believed to occur primarily during pre-GC formation (Roco et al. 2019).

A small caveat of the current model, as currently implemented on the CompuCell3D platform, is that when the cell density is too high such that the DZ is packed, there is a small chance that some dividing B cells vanish in the DZ when their volumes are too small (as represented by the small left tail of the DZ B cell volume distribution in Fig. 3H). This issue can be prevented in future iterations by imposing a limiting resource for cell growth and division to control DZ cell density. B cells are highly mobile in both the DZ and LZ with a mobility pattern observing persistent random walk (Allen et al. 2007b, Schwickert et al. 2007, Beltman et al. 2011). While we did not analyze the B cell mobility in this regard, the CompuCell3D platform, which is based on the Cellular Potts Model that follows the Boltzmann law (Graner and Glazier 1992), is capable of simulating persistent random walk (Aponte-Serrano 2021). In our case, the *temperature* parameter and local CXCL12 and 13 distribution patterns in the LZ and DZ can be optimized, along with reducing the directional chemotactic forces, to help accentuate the random-walk effect. As indicated above, it was previously showed that the persistent random walk of B cells could be an emergent behavior of B cells in a crowded GC environment (Hawkins et al. 2011), thus it will be interesting to inspect our GC model in this regard.

In the current model the two daughter cells split the parent cell’s volume by following a lognormal distribution. While this approach implemented some degree of cell division asymmetry, in future iterations asymmetrical cell divisions can be better implemented based on experimental studies which showed asymmetrical segregation of pMHCII among the two daughter cells, which plays a role in B cell fate decision-making (Thaunat et al. 2012), and was implemented in recent models (Merino Tejero et al. 2021a, Merino Tejero et al. 2021b). In the current model the length of each cell cycle in a proliferative burst is targeted as a constant. It has been shown that the S phase, which constitutes the major portion of the cell cycles of proliferating B cells, can be shortened by regulating replication fork progression, while the relative order of replication origin activation is preserved (Gitlin et al. 2015). The degree of S-phase shortening depends on the interaction strength of B cells with Tfh cells which in turn depends on BCR antigen affinity. Therefore, positively selected high-affinity GC B cells, upon returning to the DZ, will proliferate not only with a larger burst size but also with accelerated cell cycles. This could be a mechanism to compensate for the tendency of longer DZ residence time of high-affinity B cells due to more cell divisions such that they can return to the LZ sooner. Future iterations of the model may consider to incorporate affinity-dependent cell cycle shortening.

The molecular network model running in each B cell uses the Gillespie’s stochastic simulation algorithm, with some of the molecular switching actions implemented as hybrid, rule-based events. For simplicity and parsimony, several genes known to participate in GCR and B cell terminal differentiation are not included in the current implementation of the GC model, including BCL6, IRF4, BACH2, and PAX5. BCL6 is upregulated in antigen-engaged B cells in the early stage of GC formation, before these cells migrate back into the intrafollicular space and cluster in the GC (Kitano et al. 2011). There does not appear to be cyclic BCL6 expression between the DZ and LZ. Since our current model starts with B cells seeding the GC, not including the initial interactions at the T/B border, the absence of BCL6 in the molecular network should not affect the simulation results. BCL6, IRF4, BACH2, BLIMP1, and PAX5 form coupled positive or double-negative feedback loops underpinning multistability-based binary decision-making in B cells (Bhattacharya et al. 2010, Méndez and Mendoza 2016). In future iterations, the hybrid stochastic and rule-based approach of molecular network simulation can be updated by implementing relevant intracellular feedback circuits that enable bi- and multistability. To this end, single-cell RNA sequencing data of GC cells can be integrated to the molecular network model to parameterize the molecular abundance of the gene transcripts (Holmes et al. 2020).

There are a number of other considerations that can be potentially added in future iterations as well. Paracrine signals mediated by ILs and other cytokines secreted by B cells, Tfh cells and FDCs may be considered to better recapitulate the tissue-level molecular signaling milieu in the GC. Mobility of Tfh cells and their intracellular signal transduction and gen regulatory network can be included. The shapes and locations of residential CRCs and FDCs and as a result the spatial patterns of CXCL12 and 13 gradients can be fine-tuned according to immunohistochemistry data, which may ultimately affect the morphology and polarity of GCs.

### 6. Conclusions

In conclusion, we have developed a multiscale spatial computational modeling framework for GC simulation in CompuCell3D. The current model is capable of recapitulating GC features that are both qualitatively and to some extent quantitatively consistent with the literature. Given the complexity of GCR, simulations in such a modeling framework can help investigate a range of research questions on this hallmark event of high-affinity antibody production in response to viral infection and vaccination. Upon further extension and refinement, this open-source modeling framework may also help research in the area of autoimmunity and lymphoma when the GC goes awry due to genetic or environmental disruptions. Lastly, the GC modeling framework may also be utilized towards building the digital twins of the human immune system for precision medicine (Laubenbacher et al. 2022).

## Acknowledgements

This research was supported in part by NIEHS Superfund Research grant P42ES04911 and Emory Synergy grant. We would like to thank Dr. James P. Sluka for his technical assistance with CompuCell3D.

## Conflict of Interest

The authors declare that the research was conducted in the absence of any commercial or financial relationships that could be construed as a potential conflict of interest.

## Author Contributions

QZ conceived the model structure with inputs from CDS and NEK. DPM constructed and simulated the model in CompuCell3D. DPM and QZ conducted the parameter justification and estimation, and wrote the Python code for formal analysis of simulation results. DPM and QZ wrote the initial draft and revised the manuscript. CDS and NEK critically reviewed and revised the manuscript. All authors contributed to the article and approved the submitted version.

## Figure Legends

**Figure S1. (A-B)** Evolution of B cell clone sizes and clonal fractions respectively for a GC simulation resulting in single-clone dominance. **(C-D)** Evolution of B cell clone sizes and clonal fractions respectively for a GC simulation resulting in multi-clone dominance.

**Figure S2. (A-C)** Trajectory of a single B cell lineage branch that ends up in death in the DZ due to damaging BCR mutation. **(D-F)** Trajectory of a single B cell lineage branch that ends up in death in the LZ due to not being positively selected and the death timer counts down below a threshold level. Molecular and cellular variables are indicated.

## Notes

### Competing Interest Statement

The authors have declared no competing interest.

## References

Adachi, Y., T. Onodera, Y. Yamada, R. Daio, M. Tsuiji, T. Inoue, K. Kobayashi, T. Kurosaki, M. Ato and Y. Takahashi (2015). “Distinct germinal center selection at local sites shapes memory B cell response to viral escape.” Journal of Experimental Medicine 212(10): 1709–1723.

Allen, C. D., K. M. Ansel, C. Low, R. Lesley, H. Tamamura, N. Fujii and J. G. Cyster (2004). “Germinal center dark and light zone organization is mediated by CXCR4 and CXCR5.” Nat Immunol 5(9): 943–952.

Allen, C. D., T. Okada and J. G. Cyster (2007a). “Germinal-center organization and cellular dynamics.” Immunity 27(2): 190–202.

Allen, C. D., T. Okada, H. L. Tang and J. G. Cyster (2007b). “Imaging of germinal center selection events during affinity maturation.” Science 315(5811): 528–531.

Allman, D., J. R. Wilmore and B. T. Gaudette (2019). “The continuing story of T-cell independent antibodies.” Immunol Rev 288(1): 128–135.

Amitai, A., L. Mesin, G. D. Victora, M. Kardar and A. K. Chakraborty (2017). “A Population Dynamics Model for Clonal Diversity in a Germinal Center.” Front Microbiol 8: 1693.

Aponte-Serrano, J. O. (2021). Multicellular multiscale spatial modeling of the immune response to pathogens and cancer. PhD, Indiana University.

Arulraj, T., S. C. Binder and M. Meyer-Hermann (2021a). “In Silico Analysis of the Longevity and Timeline of Individual Germinal Center Reactions in a Primary Immune Response.” Cells 10(7).

Arulraj, T., S. C. Binder and M. Meyer-Hermann (2021b). “Rate of Immune Complex Cycling in Follicular Dendritic Cells Determines the Extent of Protecting Antigen Integrity and Availability to Germinal Center B Cells.” J Immunol 206(7): 1436–1442.

Arulraj, T., S. C. Binder and M. Meyer-Hermann (2022a). “Antibody Mediated Intercommunication of Germinal Centers.” Cells 11(22): 3680.

Arulraj, T., S. C. Binder and M. Meyer-Hermann (2022b). “Investigating the Mechanism of Germinal Center Shutdown.” Front Immunol 13: 922318.

Arulraj, T., S. C. Binder, P. A. Robert and M. Meyer-Hermann (2021c). “Germinal Centre Shutdown.” Frontiers in Immunology 12.

Bachmann, M. F., B. Odermatt, H. Hengartner and R. M. Zinkernagel (1996). “Induction of long-lived germinal centers associated with persisting antigen after viral infection.” J Exp Med 183(5): 2259–2269.

Bannard, O., Robert M. Horton, Christopher D. C. Allen, J. An, T. Nagasawa and Jason G. Cyster (2013). “Germinal Center Centroblasts Transition to a Centrocyte Phenotype According to a Timed Program and Depend on the Dark Zone for Effective Selection.” Immunity 39(5): 912–924.

Beltman, J. B., C. D. C. Allen, J. G. Cyster and R. J. de Boer (2011). “B cells within germinal centers migrate preferentially from dark to light zone.” Proceedings of the National Academy of Sciences 108(21): 8755–8760.

Beyer, T., M. Meyer-Hermann and G. Soff (2002). “A possible role of chemotaxis in germinal center formation.” Int Immunol 14(12): 1369–1381.

Bhagchandani, S. H., L. Yang, L. Maiorino, E. Ben-Akiva, K. A. Rodrigues, A. Romanov, H. Suh, A. Aung, S. Wu, A. Wadhera, A. K. Chakraborty and D. J. Irvine (2023). “Two-dose “extended priming” immunization amplifies humoral immune responses by synchronizing vaccine delivery with the germinal center response.” bioRxiv.

Bhattacharya, S., R. B. Conolly, N. E. Kaminski, R. S. Thomas, M. E. Andersen and Q. Zhang (2010). “A bistable switch underlying B-cell differentiation and its disruption by the environmental contaminant 2,3,7,8-tetrachlorodibenzo-p-dioxin.” Toxicol Sci 115(1): 51–65.

Binder, S. C. and M. Meyer-Hermann (2016). “Implications of Intravital Imaging of Murine Germinal Centers on the Control of B Cell Selection and Division.” Front Immunol 7: 593.

Breitfeld, D., L. Ohl, E. Kremmer, J. Ellwart, F. Sallusto, M. Lipp and R. Förster (2000). “Follicular B helper T cells express CXC chemokine receptor 5, localize to B cell follicles, and support immunoglobulin production.” J Exp Med 192(11): 1545–1552.

Calado, D. P., Y. Sasaki, S. A. Godinho, A. Pellerin, K. Köchert, B. P. Sleckman, I. M. de Alborán, M. Janz, S. Rodig and K. Rajewsky (2012). “The cell-cycle regulator c-Myc is essential for the formation and maintenance of germinal centers.” Nature Immunology 13(11): 1092–1100.

Chan, C., M. Billard, S. A. Ramirez, H. Schmidl, E. Monson and T. B. Kepler (2013). “A model for migratory B cell oscillations from receptor down-regulation induced by external chemokine fields.” Bull Math Biol 75(1): 185–205.

Choi, K., J. K. Medley, M. König, K. Stocking, L. Smith, S. Gu and H. M. Sauro (2018). “Tellurium: An extensible python-based modeling environment for systems and synthetic biology.” Biosystems 171: 74–79.

Chong, A. S. (2020). “Mechanisms of organ transplant injury mediated by B cells and antibodies: Implications for antibody-mediated rejection.” American Journal of Transplantation 20(S4): 23–32.

Chou, C., D. J. Verbaro, E. Tonc, M. Holmgren, M. Cella, M. Colonna, D. Bhattacharya and T. Egawa (2016). “The Transcription Factor AP4 Mediates Resolution of Chronic Viral Infection through Amplification of Germinal Center B Cell Responses.” Immunity 45(3): 570–582.

Cosgrove, J., M. Novkovic, S. Albrecht, N. B. Pikor, Z. Zhou, L. Onder, U. Mörbe, J. Cupovic, H. Miller, K. Alden, A. Thuery, P. O’Toole, R. Pinter, S. Jarrett, E. Taylor, D. Venetz, M. Heller, M. Uguccioni, D. F. Legler, C. J. Lacey, A. Coatesworth, W. G. Polak, T. Cupedo, B. Manoury, M. Thelen, J. V. Stein, M. Wolf, M. C. Leake, J. Timmis, B. Ludewig and M. C. Coles (2020). “B cell zone reticular cell microenvironments shape CXCL13 gradient formation.” Nature Communications 11(1): 3677.

Crotty, S. (2019). “T Follicular Helper Cell Biology: A Decade of Discovery and Diseases.” Immunity 50(5): 1132–1148.

Dominguez-Sola, D., J. Kung, A. B. Holmes, V. A. Wells, T. Mo, K. Basso and R. Dalla-Favera (2015). “The FOXO1 Transcription Factor Instructs the Germinal Center Dark Zone Program.” Immunity 43(6): 1064–1074.

Dominguez-Sola, D., G. D. Victora, C. Y. Ying, R. T. Phan, M. Saito, M. C. Nussenzweig and R. Dalla-Favera (2012). “The proto-oncogene MYC is required for selection in the germinal center and cyclic reentry.” Nature Immunology 13(11): 1083–1091.

Fagarasan, S. and T. Honjo (2000). “T-Independent immune response: new aspects of B cell biology.” Science 290(5489): 89–92.

Faili, A., S. Aoufouchi, Q. Guéranger, C. Zober, A. Léon, B. Bertocci, J. C. Weill and C. A. Reynaud (2002). “AID-dependent somatic hypermutation occurs as a DNA single-strand event in the BL2 cell line.” Nat Immunol 3(9): 815–821.

Figge, M. T., A. Garin, M. Gunzer, M. Kosco-Vilbois, K. M. Toellner and M. Meyer-Hermann (2008). “Deriving a germinal center lymphocyte migration model from two-photon data.” J Exp Med 205(13): 3019–3029.

Finkin, S., H. Hartweger, T. Y. Oliveira, E. E. Kara and M. C. Nussenzweig (2019). “Protein Amounts of the MYC Transcription Factor Determine Germinal Center B Cell Division Capacity.” Immunity 51(2): 324–336.e325.

García-Valiente, R., E. Merino Tejero, M. Stratigopoulou, D. Balashova, A. Jongejan, D. Lashgari, A. Pélissier, T. G. Caniels, M. A. F. Claireaux, A. Musters, M. J. van Gils, M. Rodríguez Martínez, N. de Vries, M. Meyer-Hermann, J. E. J. Guikema, H. Hoefsloot and A. H. C. van Kampen (2023). “Understanding repertoire sequencing data through a multiscale computational model of the germinal center.” NPJ Syst Biol Appl 9(1): 8.

Garg, A. K., R. Desikan and N. M. Dixit (2019). “Preferential Presentation of High-Affinity Immune Complexes in Germinal Centers Can Explain How Passive Immunization Improves the Humoral Response.” Cell Rep 29(12): 3946–3957.e3945.

Garg, A. K., T. Mitra, M. Schips, A. Bandyopadhyay and M. Meyer-Hermann (2023). “Amount of antigen, T follicular helper cells and affinity of founder cells shape the diversity of germinal center B cells: A computational study.” Front Immunol 14: 1080853.

Germolec, D. R., H. Lebrec, S. E. Anderson, G. R. Burleson, A. Cardenas, E. Corsini, S. E. Elmore, B. L. F. Kaplan, B. P. Lawrence, G. M. Lehmann, C. C. Maier, C. M. McHale, L. P. Myers, M. Pallardy, A. A. Rooney, L. Zeise, L. Zhang and M. T. Smith (2022). “Consensus on the Key Characteristics of Immunotoxic Agents as a Basis for Hazard Identification.” Environmental Health Perspectives 130(10): 105001.

Gillespie, D. T. (1977). “Exact stochastic simulation of coupled chemical reactions.” The Journal of Physical Chemistry 81(25): 2340–2361.

Gitlin, A. D., C. T. Mayer, T. Y. Oliveira, Z. Shulman, M. J. K. Jones, A. Koren and M. C. Nussenzweig (2015). “T cell help controls the speed of the cell cycle in germinal center B cells.” Science 349(6248): 643–646.

Gitlin, A. D., Z. Shulman and M. C. Nussenzweig (2014). “Clonal selection in the germinal centre by regulated proliferation and hypermutation.” Nature 509(7502): 637–640.

Graner, F. and J. A. Glazier (1992). “Simulation of biological cell sorting using a two-dimensional extended Potts model.” Physical Review Letters 69(13): 2013–2016.

Hawkins, J. B., M. T. Jones, P. E. Plassmann and D. A. Thorley-Lawson (2011). “Chemotaxis in densely populated tissue determines germinal center anatomy and cell motility: a new paradigm for the development of complex tissues.” PLoS One 6(12): e27650.

Heinzel, S., T. Binh Giang, A. Kan, J. M. Marchingo, B. K. Lye, L. M. Corcoran and P. D. Hodgkin (2017). “A Myc-dependent division timer complements a cell-death timer to regulate T cell and B cell responses.” Nat Immunol 18(1): 96–103.

Heise, N., N. S. De Silva, K. Silva, A. Carette, G. Simonetti, M. Pasparakis and U. Klein (2014). “Germinal center B cell maintenance and differentiation are controlled by distinct NF-κB transcription factor subunits.” J Exp Med 211(10): 2103–2118.

Hinman, R. M., J. N. Bushanam, W. A. Nichols and A. B. Satterthwaite (2007). “B Cell Receptor Signaling Down-Regulates Forkhead Box Transcription Factor Class O 1 mRNA Expression via Phosphatidylinositol 3-Kinase and Bruton’s Tyrosine Kinase.” The Journal of Immunology 178(2): 740–747.

Holmes, A. B., C. Corinaldesi, Q. Shen, R. Kumar, N. Compagno, Z. Wang, M. Nitzan, E. Grunstein, L. Pasqualucci and R. Dalla-Favera (2020). “Single-cell analysis of germinal-center B cells informs on lymphoma cell of origin and outcome.” Journal of Experimental Medicine 217(10): e20200483.

Inoue, T. and T. Kurosaki (2024). “Memory B cells.” Nature Reviews Immunology 24(1): 5–17.

Inoue, T., I. Moran, R. Shinnakasu, T. G. Phan and T. Kurosaki (2018). “Generation of memory B cells and their reactivation.” Immunol Rev 283(1): 138–149.

Ise, W., K. Fujii, K. Shiroguchi, A. Ito, K. Kometani, K. Takeda, E. Kawakami, K. Yamashita, K. Suzuki, T. Okada and T. Kurosaki (2018). “T Follicular Helper Cell-Germinal Center B Cell Interaction Strength Regulates Entry into Plasma Cell or Recycling Germinal Center Cell Fate.” Immunity 48(4): 702–715.e704.

Kasturi, S. P., I. Skountzou, R. A. Albrecht, D. Koutsonanos, T. Hua, H. I. Nakaya, R. Ravindran, S. Stewart, M. Alam, M. Kwissa, F. Villinger, N. Murthy, J. Steel, J. Jacob, R. J. Hogan, A. García-Sastre, R. Compans and B. Pulendran (2011). “Programming the magnitude and persistence of antibody responses with innate immunity.” Nature 470(7335): 543–547.

Kim, C. H., L. S. Rott, I. Clark-Lewis, D. J. Campbell, L. Wu and E. C. Butcher (2001). “Subspecialization of CXCR5+ T cells: B helper activity is focused in a germinal center-localized subset of CXCR5+ T cells.” J Exp Med 193(12): 1373–1381.

Kitano, M., S. Moriyama, Y. Ando, M. Hikida, Y. Mori, T. Kurosaki and T. Okada (2011). “Bcl6 protein expression shapes pre-germinal center B cell dynamics and follicular helper T cell heterogeneity.” Immunity 34(6): 961–972.

Kräutler, N. J., D. Suan, D. Butt, K. Bourne, J. R. Hermes, T. D. Chan, C. Sundling, W. Kaplan, P. Schofield, J. Jackson, A. Basten, D. Christ and R. Brink (2017). “Differentiation of germinal center B cells into plasma cells is initiated by high-affinity antigen and completed by Tfh cells.” J Exp Med 214(5): 1259–1267.

Kroese, F. G., A. S. Wubbena, H. G. Seijen and P. Nieuwenhuis (1987). “Germinal centers develop oligoclonally.” Eur J Immunol 17(7): 1069–1072.

Küppers, R., M. Zhao, M. L. Hansmann and K. Rajewsky (1993). “Tracing B cell development in human germinal centres by molecular analysis of single cells picked from histological sections.” Embo j 12(13): 4955–4967.

Laidlaw, B. J. and A. H. Ellebedy (2022). “The germinal centre B cell response to SARS-CoV-2.” Nature Reviews Immunology 22(1): 7–18.

Laidlaw, B. J., T. H. Schmidt, J. A. Green, C. D. Allen, T. Okada and J. G. Cyster (2017). “The Eph-related tyrosine kinase ligand Ephrin-B1 marks germinal center and memory precursor B cells.” J Exp Med 214(3): 639–649.

Lashgari, D., E. Merino Tejero, M. Meyer-Hermann, M. A. F. Claireaux, M. J. van Gils, H. C. J. Hoefsloot and A. H. C. van Kampen (2022). “From affinity selection to kinetic selection in Germinal Centre modelling.” PLoS Comput Biol 18(6): e1010168.

Laubenbacher, R., A. Niarakis, T. Helikar, G. An, B. Shapiro, R. S. Malik-Sheriff, T. J. Sego, A. Knapp, P. Macklin and J. A. Glazier (2022). “Building digital twins of the human immune system: toward a roadmap.” npj Digital Medicine 5(1): 64.

Liu, D., H. Xu, C. Shih, Z. Wan, X. Ma, W. Ma, D. Luo and H. Qi (2015). “T–B-cell entanglement and ICOSL-driven feed-forward regulation of germinal centre reaction.” Nature 517(7533): 214–218.

Luo, W., F. Weisel and M. J. Shlomchik (2018). “B Cell Receptor and CD40 Signaling Are Rewired for Synergistic Induction of the c-Myc Transcription Factor in Germinal Center B Cells.” Immunity 48(2): 313–326.e315.

Martínez, M. R., A. Corradin, U. Klein, M. J. Álvarez, G. M. Toffolo, B. di Camillo, A. Califano and G. A. Stolovitzky (2012). “Quantitative modeling of the terminal differentiation of B cells and mechanisms of lymphomagenesis.” Proc Natl Acad Sci U S A 109(7): 2672–2677.

Mayer, C. T., A. Gazumyan, E. E. Kara, A. D. Gitlin, J. Golijanin, C. Viant, J. Pai, T. Y. Oliveira, Q. Wang, A. Escolano, M. Medina-Ramirez, R. W. Sanders and M. C. Nussenzweig (2017). “The microanatomic segregation of selection by apoptosis in the germinal center.” Science 358(6360).

Méndez, A. and L. Mendoza (2016). “A Network Model to Describe the Terminal Differentiation of B Cells.” PLoS Comput Biol 12(1): e1004696.

Merino Tejero, E., D. Lashgari, R. García-Valiente, X. Gao, F. Crauste, P. A. Robert, M. Meyer-Hermann, M. R. Martínez, S. M. van Ham, J. E. J. Guikema, H. Hoefsloot and A. H. C. van Kampen (2021a). “Multiscale Modeling of Germinal Center Recapitulates the Temporal Transition From Memory B Cells to Plasma Cells Differentiation as Regulated by Antigen Affinity-Based Tfh Cell Help.” Front Immunol 11: 620716.

Merino Tejero, E., D. Lashgari, R. García-Valiente, J. He, P. A. Robert, M. Meyer-Hermann, J. E. J. Guikema, H. Hoefsloot and A. H. C. van Kampen (2021b). “Coupled Antigen and BLIMP1 Asymmetric Division With a Large Segregation Between Daughter Cells Recapitulates the Temporal Transition From Memory B Cells to Plasma Cells and a DZ-to-LZ Ratio in the Germinal Center.” Frontiers in Immunology 12: 716240.

Merino Tejero, E., Q. Mao, D. Lashgari, R. García-Valiente, P. A. Robert, M. Meyer-Hermann, M. Rodríguez Martínez, J. E. J. Guikema, H. H. C. Hoefsloot and A. H. C. van Kampen (2022). “Multi-Scale Modeling Recapitulates the Effect of Genetic Alterations Associated With Diffuse Large B-Cell Lymphoma in the Germinal Center Dynamics.” Frontiers in Systems Biology 2: 864690.

Mesin, L., J. Ersching and G. D. Victora (2016). “Germinal Center B Cell Dynamics.” Immunity 45(3): 471–482.

Methot, S. P. and J. M. Di Noia (2017). “Molecular Mechanisms of Somatic Hypermutation and Class Switch Recombination.” Adv Immunol 133: 37–87.

Meyer-Hermann, M. (2002). “A mathematical model for the germinal center morphology and affinity maturation.” J Theor Biol 216(3): 273–300.

Meyer-Hermann, M. (2014). “Overcoming the dichotomy of quantity and quality in antibody responses.” J Immunol 193(11): 5414–5419.

Meyer-Hermann, M. (2019). “Injection of Antibodies against Immunodominant Epitopes Tunes Germinal Centers to Generate Broadly Neutralizing Antibodies.” Cell Reports 29(5): 1066–1073.e1065.

Meyer-Hermann, M. (2021). “A molecular theory of germinal center B cell selection and division.” Cell Rep 36(8): 109552.

Meyer-Hermann, M. and T. Beyer (2002). “Conclusions from two model concepts on germinal center dynamics and morphology.” Dev Immunol 9(4): 203–214.

Meyer-Hermann, M. and T. Beyer (2004). “The type of seeder cells determines the efficiency of germinal center reactions.” Bull Math Biol 66(1): 125–141.

Meyer-Hermann, M., S. C. Binder, L. Mesin and G. D. Victora (2018). “Computer Simulation of Multi-Color Brainbow Staining and Clonal Evolution of B Cells in Germinal Centers.” Front Immunol 9: 2020.

Meyer-Hermann, M., M. T. Figge and K. M. Toellner (2009). “Germinal centres seen through the mathematical eye: B-cell models on the catwalk.” Trends Immunol 30(4): 157–164.

Meyer-Hermann, M., E. Mohr, N. Pelletier, Y. Zhang, G. D. Victora and K. M. Toellner (2012). “A theory of germinal center B cell selection, division, and exit.” Cell Rep 2(1): 162–174.

Meyer-Hermann, M. E. (2007). “A CONCERTED ACTION OF B CELL SELECTION MECHANISMS.” Advances in Complex Systems 10(04): 557–580.

Meyer-Hermann, M. E. and P. K. Maini (2005a). “Cutting edge: back to “one-way” germinal centers.” J Immunol 174(5): 2489–2493.

Meyer-Hermann, M. E. and P. K. Maini (2005b). “Interpreting two-photon imaging data of lymphocyte motility.” Phys Rev E Stat Nonlin Soft Matter Phys 71(6 Pt 1): 061912.

Meyer-Hermann, M. E., P. K. Maini and D. Iber (2006). “An analysis of B cell selection mechanisms in germinal centers.” Math Med Biol 23(3): 255–277.

Michida, H., H. Imoto, H. Shinohara, N. Yumoto, M. Seki, M. Umeda, T. Hayashi, I. Nikaido, T. Kasukawa, Y. Suzuki and M. Okada-Hatakeyama (2020). “The Number of Transcription Factors at an Enhancer Determines Switch-like Gene Expression.” Cell Rep 31(9): 107724.

Mintz, M. A. and J. G. Cyster (2020). “T follicular helper cells in germinal center B cell selection and lymphomagenesis.” Immunol Rev 296(1): 48–61.

Mlynarczyk, C., L. Fontán and A. Melnick (2019). “Germinal center-derived lymphomas: The darkest side of humoral immunity.” Immunol Rev 288(1): 214–239.

Molari, M., K. Eyer, J. Baudry, S. Cocco and R. Monasson (2020). “Quantitative modeling of the effect of antigen dosage on B-cell affinity distributions in maturating germinal centers.” eLife 9: e55678.

Molari, M., R. Monasson and S. Cocco (2021). “Survival probability and size of lineages in antibody affinity maturation.” Phys Rev E 103(5-1): 052413.

Nutt, S. L., P. D. Hodgkin, D. M. Tarlinton and L. M. Corcoran (2015). “The generation of antibody-secreting plasma cells.” Nature Reviews Immunology 15(3): 160–171.

Olivieri, D. N., M. Escalona and J. Faro (2013). “Software tool for 3D extraction of germinal centers.” BMC Bioinformatics 14(6): S5.

Parker, D. C. (1993). “T cell-dependent B cell activation.” Annu Rev Immunol 11: 331–360.

Phan, T. G., D. Paus, T. D. Chan, M. L. Turner, S. L. Nutt, A. Basten and R. Brink (2006). “High affinity germinal center B cells are actively selected into the plasma cell compartment.” J Exp Med 203(11): 2419–2424.

Quast, I., A. R. Dvorscek, C. Pattaroni, T. M. Steiner, C. I. McKenzie, C. Pitt, K. O’Donnell, Z. Ding, D. L. Hill, R. Brink, M. J. Robinson, D. Zotos and D. M. Tarlinton (2022). “Interleukin-21, acting beyond the immunological synapse, independently controls T follicular helper and germinal center B cells.” Immunity 55(8): 1414–1430.e1415.

Reboldi, A. and J. G. Cyster (2016). “Peyer’s patches: organizing B-cell responses at the intestinal frontier.” Immunol Rev 271(1): 230–245.

Reinhardt, R. L., H.-E. Liang and R. M. Locksley (2009). “Cytokine-secreting follicular T cells shape the antibody repertoire.” Nature Immunology 10(4): 385–393.

Reshetova, P., B. D. C. van Schaik, P. L. Klarenbeek, M. E. Doorenspleet, R. E. E. Esveldt, P.-P. Tak, J. E. J. Guikema, N. de Vries and A. H. C. van Kampen (2017). “Computational Model Reveals Limited Correlation between Germinal Center B-Cell Subclone Abundancy and Affinity: Implications for Repertoire Sequencing.” Frontiers in Immunology 8.

Robert, P. A., A. Rastogi, S. C. Binder and M. Meyer-Hermann (2017). How to Simulate a Germinal Center. Germinal Centers: Methods and Protocols. D. P. Calado. New York, NY, Springer New York: 303–334.

Roco, J. A., L. Mesin, S. C. Binder, C. Nefzger, P. Gonzalez-Figueroa, P. F. Canete, J. Ellyard, Q. Shen, P. A. Robert, J. Cappello, H. Vohra, Y. Zhang, C. R. Nowosad, A. Schiepers, L. M. Corcoran, K. M. Toellner, J. M. Polo, M. Meyer-Hermann, G. D. Victora and C. G. Vinuesa (2019). “Class-Switch Recombination Occurs Infrequently in Germinal Centers.” Immunity 51(2): 337–350.e337.

Rodda, L. B., O. Bannard, B. Ludewig, T. Nagasawa and J. G. Cyster (2015). “Phenotypic and morphological properties of germinal center dark zone Cxcl12-expressing reticular cells.” The Journal of Immunology 195(10): 4781–4791.

Roy, K., S. Mitchell, Y. Liu, S. Ohta, Y. S. Lin, M. O. Metzig, S. L. Nutt and A. Hoffmann (2019). “A Regulatory Circuit Controlling the Dynamics of NFκB cRel Transitions B Cells from Proliferation to Plasma Cell Differentiation.” Immunity 50(3): 616–628.e616.

Sander, S., Van T. Chu, T. Yasuda, A. Franklin, R. Graf, Dinis P. Calado, S. Li, K. Imami, M. Selbach, M. Di Virgilio, L. Bullinger and K. Rajewsky (2015). “PI3 Kinase and FOXO1 Transcription Factor Activity Differentially Control B Cells in the Germinal Center Light and Dark Zones.” Immunity 43(6): 1075–1086.

Schwickert, T. A., R. L. Lindquist, G. Shakhar, G. Livshits, D. Skokos, M. H. Kosco-Vilbois, M. L. Dustin and M. C. Nussenzweig (2007). “In vivo imaging of germinal centres reveals a dynamic open structure.” Nature 446(7131): 83–87.

Sharbeen, G., C. W. Yee, A. L. Smith and C. J. Jolly (2012). “Ectopic restriction of DNA repair reveals that UNG2 excises AID-induced uracils predominantly or exclusively during G1 phase.” J Exp Med 209(5): 965–974.

Shinnakasu, R., T. Inoue, K. Kometani, S. Moriyama, Y. Adachi, M. Nakayama, Y. Takahashi, H. Fukuyama, T. Okada and T. Kurosaki (2016). “Regulated selection of germinal-center cells into the memory B cell compartment.” Nature Immunology 17(7): 861–869.

Shinohara, H., M. Behar, K. Inoue, M. Hiroshima, T. Yasuda, T. Nagashima, S. Kimura, H. Sanjo, S. Maeda, N. Yumoto, S. Ki, S. Akira, Y. Sako, A. Hoffmann, T. Kurosaki and M. Okada-Hatakeyama (2014). “Positive feedback within a kinase signaling complex functions as a switch mechanism for NF-κB activation.” Science 344(6185): 760–764.

Smith, K. G. C., A. Light, G. J. V. Nossal and D. M. Tarlinton (1997). “The extent of affinity maturation differs between the memory and antibody-forming cell compartments in the primary immune response.” The EMBO Journal 16(11): 2996–3006.

Stebegg, M., S. D. Kumar, A. Silva-Cayetano, V. R. Fonseca, M. A. Linterman and L. Graca (2018). “Regulation of the Germinal Center Response.” Frontiers in Immunology 9.

Stewart, I., D. Radtke, B. Phillips, S. J. McGowan and O. Bannard (2018). “Germinal Center B Cells Replace Their Antigen Receptors in Dark Zones and Fail Light Zone Entry when Immunoglobulin Gene Mutations are Damaging.” Immunity 49(3): 477–489.e477.

Swat, M. H., G. L. Thomas, J. M. Belmonte, A. Shirinifard, D. Hmeljak and J. A. Glazier (2012). Multi-Scale Modeling of Tissues Using CompuCell3D. Methods in Cell Biology. A. R. Asthagiri and A. P. Arkin, Academic Press. 110: 325–366.

Tas, J. M., L. Mesin, G. Pasqual, S. Targ, J. T. Jacobsen, Y. M. Mano, C. S. Chen, J. C. Weill, C. A. Reynaud, E. P. Browne, M. Meyer-Hermann and G. D. Victora (2016). “Visualizing antibody affinity maturation in germinal centers.” Science 351(6277): 1048–1054.

Thaunat, O., A. G. Granja, P. Barral, A. Filby, B. Montaner, L. Collinson, N. Martinez-Martin, N. E. Harwood, A. Bruckbauer and F. D. Batista (2012). “Asymmetric segregation of polarized antigen on B cell division shapes presentation capacity.” Science 335(6067): 475–479.

Vaidehi Narayanan, H. and A. Hoffmann (2022). “From Antibody Repertoires to Cell-Cell Interactions to Molecular Networks: Bridging Scales in the Germinal Center.” Frontiers in Immunology 13.

Valeri, V., A. Sochon, C. Ye, X. Mao, D. Lecoeuche, S. Fillatreau, J.-C. Weill, C.-A. Reynaud and Y. Hao (2022). “B cell intrinsic and extrinsic factors impacting memory recall responses to SRBC challenge.” Frontiers in Immunology 13.

Verstegen, N. J. M., V. Ubels, H. V. Westerhoff, S. M. van Ham and M. Barberis (2021). “System-Level Scenarios for the Elucidation of T Cell-Mediated Germinal Center B Cell Differentiation.” Frontiers in Immunology 12.

Viant, C., G. H. J. Weymar, A. Escolano, S. Chen, H. Hartweger, M. Cipolla, A. Gazumyan and M. C. Nussenzweig (2020). “Antibody Affinity Shapes the Choice between Memory and Germinal Center B Cell Fates.” Cell 183(5): 1298–1311.e1211.

Victora, G. D., D. Dominguez-Sola, A. B. Holmes, S. Deroubaix, R. Dalla-Favera and M. C. Nussenzweig (2012). “Identification of human germinal center light and dark zone cells and their relationship to human B-cell lymphomas.” Blood 120(11): 2240–2248.

Victora, G. D. and M. C. Nussenzweig (2022). “Germinal Centers.” Annual Review of Immunology 40(1): 413–442.

Victora, G. D., T. A. Schwickert, D. R. Fooksman, A. O. Kamphorst, M. Meyer-Hermann, M. L. Dustin and M. C. Nussenzweig (2010). “Germinal center dynamics revealed by multiphoton microscopy with a photoactivatable fluorescent reporter.” Cell 143(4): 592–605.

Vinuesa, C. G., M. A. Linterman, C. C. Goodnow and K. L. Randall (2010). “T cells and follicular dendritic cells in germinal center B-cell formation and selection.” Immunol Rev 237(1): 72–89.

Wang, P., C. M. Shih, H. Qi and Y. H. Lan (2016). “A Stochastic Model of the Germinal Center Integrating Local Antigen Competition, Individualistic T-B Interactions, and B Cell Receptor Signaling.” J Immunol 197(4): 1169–1182.

Wang, Q., K.-R. Kieffer-Kwon, T. Y. Oliveira, C. T. Mayer, K. Yao, J. Pai, Z. Cao, M. Dose, R. Casellas, M. Jankovic, M. C. Nussenzweig and D. F. Robbiani (2017). “The cell cycle restricts activation-induced cytidine deaminase activity to early G1.” Journal of Experimental Medicine 214(1): 49–58.

Wang, S. (2017). “Optimal Sequential Immunization Can Focus Antibody Responses against Diversity Loss and Distraction.” PLoS Comput Biol 13(1): e1005336.

Wang, S., J. Mata-Fink, B. Kriegsman, M. Hanson, D. J. Irvine, H. N. Eisen, D. R. Burton, K. D. Wittrup, M. Kardar and A. K. Chakraborty (2015). “Manipulating the selection forces during affinity maturation to generate cross-reactive HIV antibodies.” Cell 160(4): 785–797.

Wang, X., B. Cho, K. Suzuki, Y. Xu, J. A. Green, J. An and J. G. Cyster (2011). “Follicular dendritic cells help establish follicle identity and promote B cell retention in germinal centers.” Journal of Experimental Medicine 208(12): 2497–2510.

Weber, T. S. (2018). “Cell Cycle-Associated CXCR4 Expression in Germinal Center B Cells and Its Implications on Affinity Maturation.” Frontiers in Immunology 9.

Wibisana, J. N., T. Inaba, H. Shinohara, N. Yumoto, T. Hayashi, M. Umeda, M. Ebisawa, I. Nikaido, Y. Sako and M. Okada (2022). “Enhanced transcriptional heterogeneity mediated by NF-κB super-enhancers.” PLoS Genet 18(6): e1010235.

Wittenbrink, N., T. S. Weber, A. Klein, A. A. Weiser, W. Zuschratter, M. Sibila, J. Schuchhardt and M. Or-Guil (2010). “Broad volume distributions indicate nonsynchronized growth and suggest sudden collapses of germinal center B cell populations.” J Immunol 184(3): 1339–1347.

Woods, M., Y. R. Zou and A. Davidson (2015). “Defects in Germinal Center Selection in SLE.” Front Immunol 6: 425.

Xin, G., R. Zander, D. M. Schauder, Y. Chen, J. S. Weinstein, W. R. Drobyski, V. Tarakanova, J. Craft and W. Cui (2018). “Single-cell RNA sequencing unveils an IL-10-producing helper subset that sustains humoral immunity during persistent infection.” Nature Communications 9(1): 5037.

Yan, Z., H. Qi and Y. Lan (2022). “The role of geometric features in a germinal center.” Mathematical Biosciences and Engineering 19(8): 8304–8333.

Yang, L., M. Van Beek, Z. Wang, F. Muecksch, M. Canis, T. Hatziioannou, P. D. Bieniasz, M. C. Nussenzweig and A. K. Chakraborty (2023). “Antigen presentation dynamics shape the antibody response to variants like SARS-CoV-2 Omicron after multiple vaccinations with the original strain.” Cell Rep 42(4): 112256.

Young, C. and R. Brink (2021). “The unique biology of germinal center B cells.” Immunity 54(8): 1652–1664.

Zarnegar, B., J. Q. He, G. Oganesyan, A. Hoffmann, D. Baltimore and G. Cheng (2004). “Unique CD40-mediated biological program in B cell activation requires both type 1 and type 2 NF-kappaB activation pathways.” Proc Natl Acad Sci U S A 101(21): 8108–8113.

Zhang, J. and E. I. Shakhnovich (2010). “Optimality of mutation and selection in germinal centers.” PLoS Comput Biol 6(6): e1000800.

Zhang, Y., M. Meyer-Hermann, L. A. George, M. T. Figge, M. Khan, M. Goodall, S. P. Young, A. Reynolds, F. Falciani, A. Waisman, C. A. Notley, M. R. Ehrenstein, M. Kosco-Vilbois and K. M. Toellner (2013). “Germinal center B cells govern their own fate via antibody feedback.” J Exp Med 210(3): 457–464.

