## Supplementary figures and images for "A Multiscale Spatial Modeling Framework for the Germinal Center Response"

### Supplemental Figures

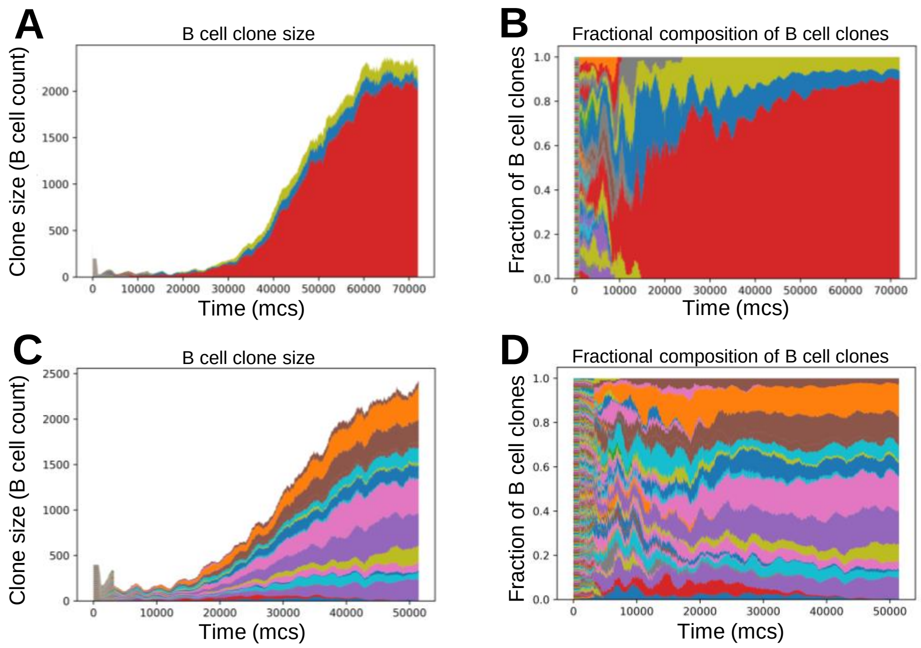

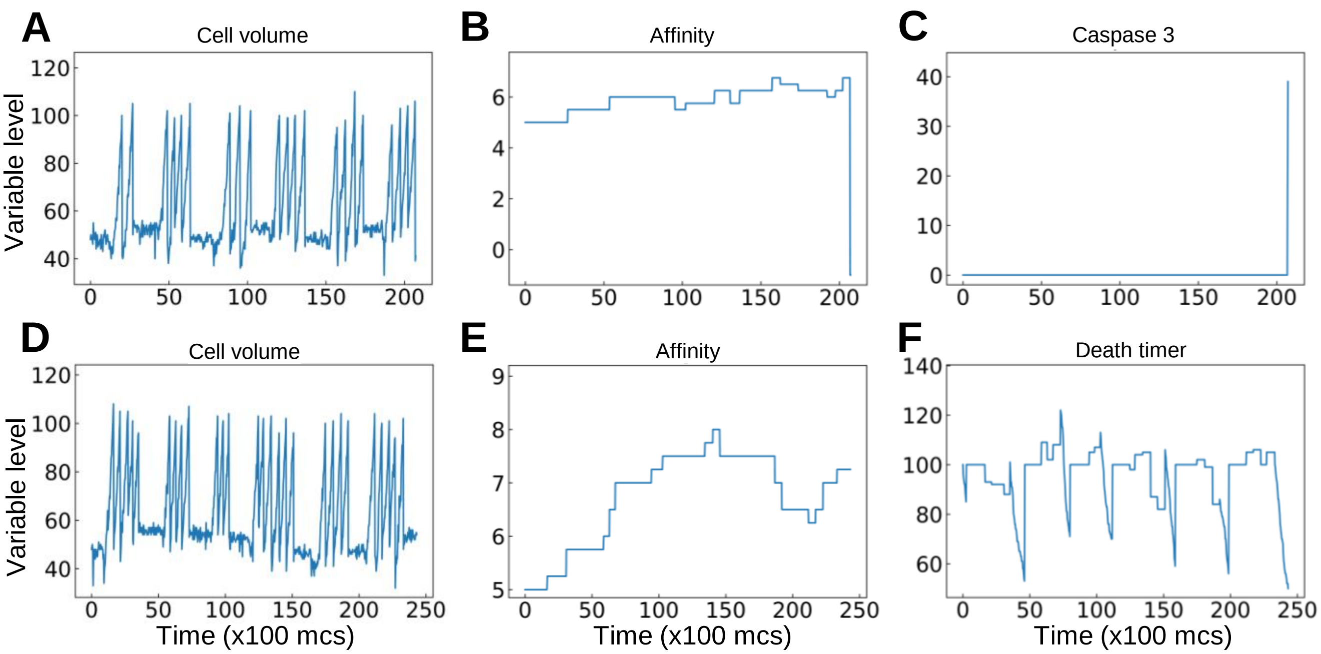
